# Urbanization and a green corridor do not impact genetic divergence in common milkweed (*Asclepias syriaca*)

**DOI:** 10.1101/2023.04.04.535613

**Authors:** Sophie T. Breitbart, Anurag A. Agrawal, Helene H. Wagner, Marc T.J. Johnson

**Affiliations:** Department of Ecology and Evolutionary Biology University of Toronto 25 Willcocks Street Toronto, Ontario Canada M5S 3B2; Department of Biology University of Toronto Mississauga 3359 Mississauga Road Mississauga, ON Canada L5L 1C6; Centre for Urban Environments University of Toronto Mississauga 3359 Mississauga Road Mississauga, ON Canada L5L 1C6; Department of Ecology & Evolutionary Biology Cornell University E145 Corson Hall Ithaca, NY USA 14853; Department of Entomology Cornell University 2126 Comstock Hall Ithaca, NY USA 14853

## Abstract

Urbanization is altering landscapes globally at an unprecedented rate. While ecological differences between urban and rural environments often promote phenotypic divergence among populations, it is unclear to what degree these trait differences arise from genetic divergence as opposed to phenotypic plasticity. Furthermore, little is known about how specific landscape elements, such as green corridors, impact genetic divergence in urban environments. We tested the hypotheses that: 1) urbanization, and 2) proximity to an urban green corridor influence genetic divergence in *Asclepias syriaca* (common milkweed) populations for phenotypic traits. Using seeds from 52 populations along three urban-to-rural subtransects in the Greater Toronto Area, one of which followed a green corridor, we grew ∼1000 plants in a common garden and observed >20 ecologically-important traits associated with plant defense/damage, reproduction, and growth over four years. We found significant heritable variation for eight traits within *A. syriaca* populations and weak phenotypic divergence among populations. However, neither urbanization nor an urban green corridor influenced genetic divergence in individual traits or multivariate phenotype. These findings contrast with the expanding literature demonstrating that urbanization promotes rapid evolutionary change and offer preliminary insights into the eco- evolutionary role of green corridors in urban environments.

## Introduction

The intense environmental changes associated with urbanization can profoundly alter the ecological processes of urban-dwelling populations. Several biotic and abiotic factors differentiate urban from non-urban environments including higher pollution, habitat fragmentation, and temperature due to the urban heat island effect^1, 2^. In turn, these alterations can impact populations and communities directly by affecting species richness^3, 4^, abundance^5, 6^, and spatiotemporal distribution^7, 8^. These environmental conditions can also impose indirect impacts by restructuring species interactions such as antagonisms (e.g., predator/prey, including herbivory) and mutualisms (e.g., plant-pollinator and plant-microbe)^9–11^. In response, individuals and ultimately populations have experienced trait changes associated with growth, reproduction, defense/damage, morphology, and behavior, a process indicative of intraspecific phenotypic divergence^12–16^. Despite the recent rise of studies documenting phenotypic divergence among urban and rural populations, few have distinguished the degree to which evolution and phenotypic plasticity drive these patterns.

Phenotypic divergence can be explained by two general mechanisms: phenotypic plasticity and genetic divergence. Phenotypic plasticity occurs when individuals with identical genotypes exhibit different phenotypes in different environments^17^ and was shown to account for the majority of the increased heat tolerance exhibited by urban water fleas (*Daphnia magna*) in a common garden experiment^18^. Conversely, genetic divergence happens via evolution and is, in urban environments, most commonly attributed to changes in natural selection, genetic drift, and gene flow^19–21^. Specifically, the widespread habitat fragmentation characteristic of urban environments is frequently implicated in reports of heightened genetic drift within populations^22, 23^, restricted gene flow among populations^24, 25^, and natural selection^26^.

Several studies have investigated the basis of phenotypic divergence in plants^27–30^, yet few include multiple traits, or traits from diverse ecological functions such as reproduction, defense/damage, and growth. Thus, we have a poor understanding of how urbanization affects divergence in the multivariate phenotype^but^ ^see^ ^16^. Individual responses to environmental change often involve multiple correlated traits^31, 32^, and excluding traits from major categories (e.g., defense) complicates the detection of shifts in life history strategies, including trade-offs, or the lack thereof. Thus, surveying a broad range of phenotypic traits is essential for determining the drivers of phenotypic divergence in urban environments.

Individual elements of the urban landscape are thought to impact the evolution of phenotypic traits. Long, connected habitat patches—henceforth, green corridors— have been shown to impact evolution in non-urban areas^33–35^ such as forest fragments, frequently by facilitating gene flow^36, 37^ and alleviating effects of genetic drift^38^, ^but^ ^see^ ^39^. In turn, these processes can accelerate adaptation through the introduction of beneficial alleles or slow it through the introduction of deleterious or neutral alleles^40–43^. However, virtually nothing is known about how these corridors impact phenotypic trait evolution in urban areas. Broadly, urban green corridors frequently impact dispersal and gene flow^44, 45^, such as in the white-footed mouse (*Peromyscus leucopus*) of New York where vegetation corridors facilitate dispersal^46^. Investigating how urban green corridors influence genetic divergence in phenotypic traits is essential for discerning whether aspects of urban landscapes shape the phenotype of urban-dwelling taxa. This knowledge would also offer conservation agencies important context about the evolutionary consequences of corridors primarily utilized to remedy the ecological costs of habitat fragmentation in urban environments^47^. Evaluating how urban green corridors impact the evolution of phenotypic traits represents an important step towards understanding evolutionary dynamics in heterogeneous urban landscapes.

Prior research demonstrating how urbanization influences ecological change in a native plant of conservation importance (*Asclepias syriaca*), as well as its pollinators and herbivores, suggests that evolutionary change may follow. In an observational survey, urbanization influenced reproductive success and pollinator community structure, while in urban populations, proximity to a green corridor inconsistently affected reproductive success, but uniformly decreased pollinator diversity and richness^48^. Likewise, another study highlighted the complexity with which nine specialist herbivore species were impacted by urbanization, season, and surveyed city^49^. Urban *A. syriaca* populations showed higher herbivore species richness yet less leaf herbivory than rural populations, with these effects varying by season and city. Additionally, the leaf-mining fly (*Liriomyza asclepiadis*) was 90% more abundant in urban areas and the milkweed stem weevil (*Rhyssomatus lineaticollis*) was 41% more abundant in rural areas, though the effect for *R. lineaticollis* varied by city. Consequently, differential ecological stressors within urban areas could drive the evolution of defense and reproductive traits in *A. syriaca* in particular.

Here, we used a common garden experiment to test the hypotheses that: 1) urbanization, and 2) proximity to an urban green corridor drive genetic divergence in phenotypic traits among populations of *A. syriaca*. We asked three questions: Q1) Is there heritable variation for phenotypic traits within populations and genetic divergence between populations? These are two necessary conditions if *A. syriaca* is to evolve in response to urban environmental change. Q2) To what extent is urbanization associated with genetic divergence in phenotypic traits among populations? If *A. syriaca* has experienced genetic divergence in response to urbanization, then population-level phenotypic trait estimates will correlate with local urbanization levels. Q3) Is proximity to an urban green corridor related to the levels of genetic divergence among urban populations? If so, there will be distinct differences in phenotypic trait estimates between populations near to and far from the corridor. This study builds upon work investigating how urbanization and a green corridor influence phenotypic divergence in reproduction in *A. syriaca*^48^ by uncovering the extent to which evolution and phenotypic plasticity shape phenotypic divergence in reproductive, growth, and defense-related traits. These results provide insights into how plants respond to rapid environmental change in heterogeneous urban environments.

## Methods

### Study system

Common milkweed, *Asclepias syriaca L.*, is an herbaceous perennial plant native to eastern North America. Though *A. syriaca* grows in discrete patches of one to thousands of ramets, often in old rural fields^50^, urban populations tend to be smaller and inhabit public parks, railway and transmission rights-of-way, and roadsides, as well as private lawns and gardens. Plants can reproduce vegetatively through rhizomes that generate clonal ramets, or sexually through the cross-pollination of hermaphroditic flowers, each with five pollen sac pairs collectively called pollinaria^50–52^. Fertilization following successful pollination by insects, such as *Apis mellifera*, *Bombus* spp., and *Halictidae* spp. in urban areas^48, 53, 54^, yields follicles (i.e., fruits) filled with wind-dispersed seeds^51, 55^.

Multiple traits protect *A. syriaca* against herbivores. Milkweeds contain a pressurized, milky sap called latex that can physically interfere with herbivores chewing tissue by gumming their mouthparts^56–58^. Herbivores that can overcome this barrier must also tolerate cardenolides, a suite of secondary metabolites which can disrupt sodium-potassium pumps (Na^+^/K^+^-ATPases) required for maintaining membrane potential^59^. At least 16 herbivores have coevolved with *Asclepias* and developed tolerance to these defenses^59, 60^, and at least nine, including the Monarch butterfly (*Danaus plexippus*), can enhance their own toxicities by sequestering cardenolides^60, 61^. Compared to growth and reproduction, heritabilities of defense traits (e.g., latex and cardenolides) are often high^62–66^, indicating an increased likelihood for traits in this category to evolve.

### Field sampling of maternal families

To evaluate whether urbanization affects the evolution of ecologically important phenotypic traits, we sampled plants at 52 sites along a 67 km urban–rural gradient in the Greater Toronto Area, Ontario, Canada (Fig. 1) as described in Breitbart et al.^48^. This gradient was split into two parallel urban subtransects ca. 5 km apart, and a third rural subtransect, all approximately the same length. The northern urban subtransect (“Urban: Non-Corridor”, N = 17 sites) crossed through the urban matrix while the southern urban subtransect (“Urban: Corridor”, N = 19 sites) followed a green corridor: a mostly continuous vegetated strip of land, ca. 50–350 m wide, that ran adjacent to linear features like trails, rights-of-way, and railways. Sampling sites were located within or near the corridor. The rural subtransect (“Rural”, N = 16 sites) extended westward from the suburban termini of the urban subtransects. Sampling sites (henceforth “populations”) of isolated patches of *A. syriaca* were spaced >500 m apart. We then divided the population into five equally-sized sections and collected seeds from one follicle (i.e., full-sibling seed family) per ramet per section, aiming to separate ramets by >3 m between ramets to avoid resampling the same clone. Seeds were stored in coin envelopes and preserved at -20°C until germination.

**Figure 1.**
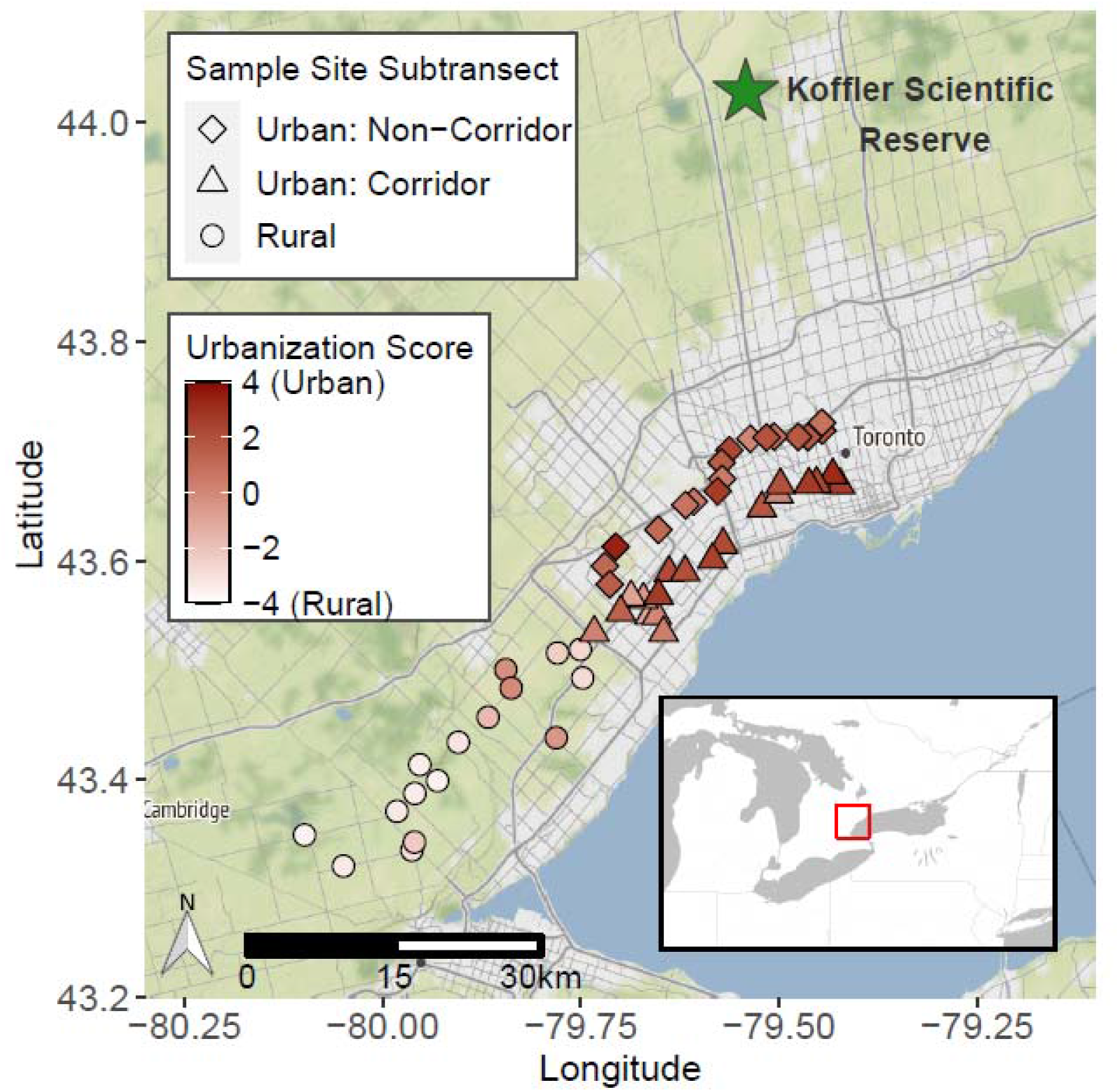
Map of 52 common milkweed populations sampled along Toronto’s urban-rural gradient and location of common garden experiment. Urban: Non-Corridor populations (squares; NLJ=LJ17). Urban: Corridor populations (triangles; NLJ=LJ19). Rural populations (circles; NLJ=LJ16). The color of the symbols indicates urbanization score, where positive values indicate a high degree of urbanization (based on the quantity of vegetation, buildings, and paved roads per 1 km²). The Stamen terrain basemap shows urban and suburban areas in light gray, nonurban agricultural and forested areas in green, and Lake Ontario in blue. Map tiles by Stamen Design, under CC BY 3.0. Data by OpenStreetMap, under ODbL.

### Urbanization metrics

We quantified urbanization using two methods as described in Breitbart et al.^48^. Briefly, we first measured the distance from each population to the Toronto urban center (43.6563, -79.3809), as a proxy for the degree of urbanization. Distance from the urban center is correlated with numerous environmental factors associated with urbanization^67^, and, for this city specifically, has been shown to influence the ecology and evolution of other plant, herbivore, and pollinator communities^10, 68, 69^. We also calculated an urbanization score for each population with the UrbanizationScore software^70–72^.

### Common garden experiment

To test if urbanization and an urban green corridor drive genetic divergence in phenotypic traits, we conducted a common garden experiment from 2019-2022. We germinated ∼10 full-sibling seeds from each of 5 families per population, then grew seeds in pots in a growth chamber for six weeks. Seedlings were then snipped at the base of the ramet to ensure equal baseline heights and transported to the University of Toronto’s Koffler Scientific Reserve (44.029, -79.531) in King City, Ontario, Canada (http://ksr.utoronto.ca/) in May 2019. Seedlings were transplanted into 3.79 L circular pots filled with field soil and topped with 2.5 cm Triple Mix, then randomized into 4 contiguous spatial blocks with 1.5 m spacing between rows and columns (Supplementary Fig. S1). Each row was covered with a 1 m-wide strip of landscaping fabric (Quest Better Barriers, Brampton, Canada) to prevent weeds from outcompeting experimental plants. Overall, we planted 954 individuals from 52 populations, 208 full-sib families, and 3 subtransects (mean: 4 families/population; 4.55 plants/family; 18.33 plants/population; range: 1-25 plants/population), with variation in representation due to germination success. We measured ecologically important traits (described below) related to plant growth, defense/damage, and reproduction over four years.

### Trait measurements

In total, we measured 21 traits from 3 functional categories: plant defense/damage, growth/development, and reproduction, as well as herbivore abundance to contextualize defense trait expression (Table 1, Supplementary Fig. S2). Measuring this large number of traits with diverse ecological functions allowed us to assess how the multivariate phenotype of *A. syriaca* was evolving in response to urban environmental gradients.

**Table 1.** Broad-sense heritability (H^2^), coefficient of genetic variation (CV_G_), heritable genetic variation within populations (p (Family)), standardized measure of genetic differentiation among populations for quantitative traits (Q_ST_), and heritable genetic variation among populations (p (Population)) for phenotypic traits.

| | $H^2$ | $CV_G$ | $p$<br>(Family) | $Q_{ST}$ | $p$<br>(Population) |
| --- | --- | --- | --- | --- | --- |
| *Defense/Damage* |  |  |  |  |  |
| Latex exudation | 0.092 | 0.015 | 0.077 | 0.174 | **0.016** |
| Herbivory before flowering (binary) | 0.021 | 0.271 | 0.490 | 0.000 | 0.500 |
| Herbivory before flowering (quantitative) | 0.000 | 0.000 | 0.500 | 1.000 | 0.500 |
| Herbivory after flowering (binary) | 0.588 | 0.948 | **0.004** | 0.000 | 0.496 |
| Herbivory after flowering (quantitative) | 0.043 | 4.456 | 0.260 | 0.000 | 0.500 |
| Weevil damage (binary) | 0.418 | 0.995 | **0.030** | 0.000 | 0.500 |
| Weevil damage (quantitative) | 0.080 | 0.031 | 0.138 | 0.116 | 0.136 |
| *Reproduction* |  |  |  |  |  |
| Flowering success | 0.102 | 0.309 | 0.302 | 0.492 | **0.014** |
| Flowers per inflorescence | 0.000 | 0.000 | 0.500 | 1.000 | 0.180 |
| Flower size | 0.082 | 0.042 | 0.370 | 0.305 | 0.134 |
| Flowering duration | 0.000 | 0.000 | 0.500 | 0.555 | 0.500 |
| Date of first flower | 0.000 | 0.010 | 0.500 | 0.000 | 0.500 |
| Pollinaria removed | 0.544 | 0.299 | **0.034** | 0.009 | 0.449 |
| Follicles | 0.000 | 0.000 | 0.500 | 0.287 | 0.500 |
| Date of first follicle | 0.235 | 0.034 | **<0.001** | 0.002 | 0.500 |
| Inflorescences | 0.000 | 0.000 | 0.496 | 1.000 | **0.035** |
| *Herbivore abundance* |  |  |  |  |  |
| *Danaus plexippus* abundance | 0.003 | 0.767 | **0.021** | 0.052 | 0.396 |
| *Liriomyza asclepiadis* abundance | 0.055 | 0.140 | 0.142 | 0.162 | 0.106 |
| *Labidomera clivicollis* abundance | 0.000 | 0.011 | 0.500 | 0.122 | 0.500 |
| *Growth* |  |  |  |  |  |
| LDMC | 0.000 | 0.000 | 0.500 | 0.011 | 0.380 |
| SLA | 0.000 | 0.000 | 0.500 | 1.000 | 0.404 |
| Height before flowering | 0.069 | 0.004 | 0.112 | 0.000 | 0.500 |
| Height after flowering | 0.132 | 0.008 | **0.009** | 0.052 | 0.189 |
| Relative growth rate | 0.021 | 3.643 | 0.394 | 0.000 | 0.500 |
| Ramets before flowering | 0.161 | 0.127 | **<0.001** | 0.009 | 0.429 |
| Ramets after flowering | 0.138 | 0.189 | **<0.001** | 0.017 | 0.364 |
| Mortality | 0.178 | 0.320 | 0.189 | 0.139 | 0.186 |

We assessed five ecologically important traits related to plant defense/damage (Supplementary Text S1). We measured leaf herbivory by chewing herbivores before and after flowering by selecting the five oldest leaves pointing east and visually estimating the percent leaf area removed as described in Johnson et al.^73^. We quantified stem damage by a specialist milkweed stem weevil, *Rhyssomatus lineaticollis* (Say, 1824), as the sum length of the oviposition scars per plant, which strongly predicts the number of eggs deposited into the ramet^74^. Latex exudation was quantified by snipping the tip (∼0.5 cm) off the youngest fully expanded intact leaf and collecting the latex exudate on a pre-weighed 1cm filter paper disc as described in Agrawal et al.^62^. Discs were placed in a pre-weighed microcentrifuge tube on dry ice, then transferred to a -80°C freezer until weighed to the nearest 1 µg.

We generated population-level estimates of leaf cardenolide concentrations by freeze-drying the leaf used to assess latex exudation and its opposite at -80°C, then finely grinding 50 mg of pooled tissue per population and following the protocol described in Petschenka et al.^75^.

We assessed the abundance of the ten mostly host-specific herbivore species. In mid-July and mid-August 2020, we surveyed the abundance of ten specialist herbivore species located on all aboveground parts of the plants (Supplementary Fig. S2). The total number of leaf splotch mines represented the abundance of *Liriomyza asclepiadis.* Monarch butterfly (*Danaus plexippus*) eggs, caterpillars, and chrysalises all contributed towards abundance. In 2021, we repeated the surveys for three species: *Danaus plexippus, Liriomyza asclepiadis* and *Labidomera clivicollis*.

We assessed eight traits associated with plant growth and development. We measured the height of all ramets before and after flowering began as the length of the ramet from soil level to the apical meristem, a surrogate of ramet biomass^48^. Mortality was recorded in early September, and we counted the number of ramets per pot before and after flowering began as each single shoot emerging from the soil or a shoot connected to another shoot at the soil surface. We calculated relative growth rate as:

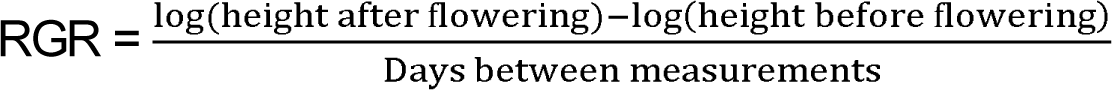

To assess specific leaf area (SLA) and leaf dry matter content (LDMC), we collected the youngest fully expanded intact leaf and measured wet mass, dry mass, length, and width. SLA and LDMC were calculated as leaf area/dry mass (m²/g) and dry mass/wet mass, respectively.

We assessed nine traits associated with plant reproduction. Flowering success represented whether a plant produced any viable, fully reflexed flowers. We recorded flowering duration as the number of days between the dates of the first fully reflexed flowers on the oldest and youngest inflorescences. For the three oldest inflorescences per plant, we recorded the number of flowers per inflorescence once at least half the flowers in an inflorescence became fully reflexed. From three flowers per inflorescence, we assessed the number of pollinaria removed and flower size. Flower size was measured as the length and width of each flower’s corolla, hood, and distance from one hood tip to the opposite side of the reproductive whorl (Supplementary Fig. S3). We counted the total number of inflorescences produced throughout the entire growing season. We recorded the date of the first mature follicle per plant when it exceeded 5.08 cm (2 in) in length as smaller follicles are the most likely to be aborted^55^, as well as the total number of mature follicles per plant.

### Statistical analyses

#### Quantitative genetic parameters

All analyses described below were performed using R v.4.1.1^76^.

We calculated three quantitative genetic parameters to understand how urbanization could affect divergence between populations and genetic variation within populations. To estimate the variances explained by population and family for each trait, we used the *glmmTMB* package^77^ (version 1.1.2.2) to fit the following general linear models using restricted maximum likelihood:

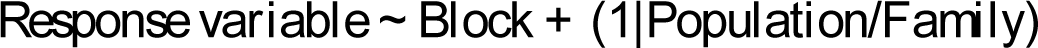

We extracted population and family-level variances with the “VarCorr” function from the *lme4* package^78^ (version 1.1.27.1), and residual variances with the “sigma” function from the *stats* package^76^ (version 4.1.1).

We calculated estimates of full-sibling broad-sense heritability (*H*²), the ratio of genetic variance to total phenotypic variance^17^, for each phenotypic trait as:

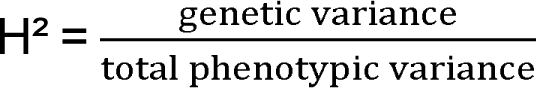

where genetic variance for full-sibs was calculated as (family-level variance)X2 and total phenotypic variance was calculated as family-level variance + population-level variance + residual variance. We calculated Q_ST_, a standardized measure of the genetic differentiation of a quantitative trait among populations^79^, as:

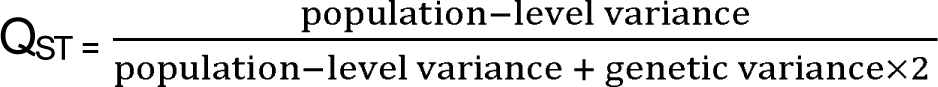

We calculated the coefficient of genetic variation (CV_G_), a dimensionless measure of evolutionary potential that is closely related to the coefficient of additive genetic variation (CV_A_)^80^, as:

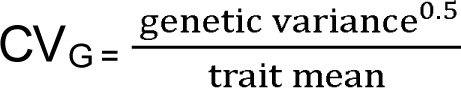

#### Genetic variation within and between populations (Q1)

To quantify heritable genetic variation for phenotypic traits within and between populations, we fitted linear mixed effect models. For response variables with Gaussian distributions, we used the “lmer” function from *lme4* to fit the following general linear mixed models using restricted maximum likelihood:

*Response ∼ Block + (1|Population/Family)*

In these models, data was restricted to the last year of measurement to minimize impacts of maternal effects. Block was treated as a fixed effect while Population and Family were random effects, with Family nested within Population. We tested the significance of Population and Family using the “ranova” function from the *lmerTest* package^81^ (version 3.1.3), then divided p-values by 2 because these were 1-sided tests (i.e., variance ≥ 0). We then obtained percent variance explained (PVE) for each random effect after extracting variances with *lme4* and calculating:

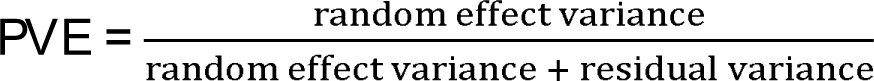

For response variables with non-Gaussian distributions, we fitted generalized linear mixed effect models with the same formula using the “glmer” function from *lme4*. To test the significance of Population and Family, we used the “PBmodcomp” function from the *pbkrtest* package^82^ (version 0.5.1) with 10,000 simulations, then divided p-values by 2 because these were 1-sided tests (i.e., variance ≥ 0). Results for ten models analyzed with 10,000 simulations were identical to those analyzed with 10, so we used 10 simulations for the remaining models. We refitted models to Gaussian distributions, then extracted variances and calculated PVE for each random effect as described above. We inspected model diagnostics with the *DHARMa*^83^ (version 0.4.3) and *performance*^84^ (version 0.7.3) packages and transformed response variables to meet the assumptions of normality and homogeneity of variance when necessary (Supplementary Tables S1-2). Response variables associated with plant herbivory and weevil damage were analyzed with manual hurdle models^85^ to account for an excess of zeroes and evaluate if herbivory/damage was present and, if so, the quantity of damage.

#### Genetic clines along an urban-rural gradient (Q2)

To test if urbanization caused genetic divergence among populations, we fitted linear mixed effect models with urbanization added as a fixed effect using the same packages described above. For both general and generalized linear mixed effect models, urbanization was added to models as both Distance to the City Center and Urbanization Score, separately. To test the significance of random effects (i.e., Population & Family), we repeated the remaining steps as described for Question 1 above. If urbanization explained heritable genetic variation within and/or between populations (i.e., p-values for Question 1 models were significant for Family and/or Population), then these p-values should have been higher in Question 2 models’ results. To test the significance of fixed effects (i.e., urbanization), we adjusted the models to use maximum likelihood and then computed ^2^ and p-values from a type II sums-of-squares ANOVA with the *car* package^86^ (version 3.0.11). If urbanization explained genetic variation within and/or between populations, then some variance should have shifted from Population and/or Family to urbanization and the p-values for urbanization should have become significant.

To test for differences associated with urbanization in overall phenotype, as opposed to differences in individual traits that could reflect specific selection pressures, we performed a multivariate phenotype analysis. This analysis incorporated all traits in the dataset and accounted for non-independence between correlated traits. We computed best linear unbiased predictions (BLUPs) for each model with the “ranef” function from *lme4*, placed these values in a response matrix, and then used the *mvabund* package^87^ (version 4.2.1) to fit general linear models and examine how multivariate phenotype varied with urbanization (both Distance and Urbanization Score) using one-way ANOVA.

#### Genetic differentiation between a green corridor & urban matrix (Q3)

To evaluate how genetic variation was associated with a green corridor between urban populations (“Urban: Corridor” and “Urban: Non-Corridor”), we added Subtransect and Urbanization:Subtransect interaction terms to the linear mixed effect models described for Question 2. For both general and generalized linear mixed effect models, we tested the significance of the random effects (i.e., Population & Family) by repeating the steps as described for Question 2 above. To test the significance of the fixed effects (i.e., Urbanization, Subtransect, and their interaction), we fitted reduced models without the interaction term, ranked full and reduced models based on Akaike’s information criterion (AIC)^88^, and selected the model with the lowest AIC as the best model. We adjusted the models to use maximum likelihood and then computed ^2^ and p-values from a type III sums-of-squares ANOVA if the best model contained an interaction with p ≤ 0.1; otherwise, we reran the analyses with type II sums-of-squares.

We also fitted models with multiple years of data and found that the effects of urbanization and a green corridor were qualitatively identical to those reported in Tables 2-3 in 85% of cases (Supplementary Text, Supplementary Table S3). We then repeated the multivariate analyses for these models as described for Question 2. Cardenolides were not analyzed due to low sample size among urban populations.

**Table 2.**
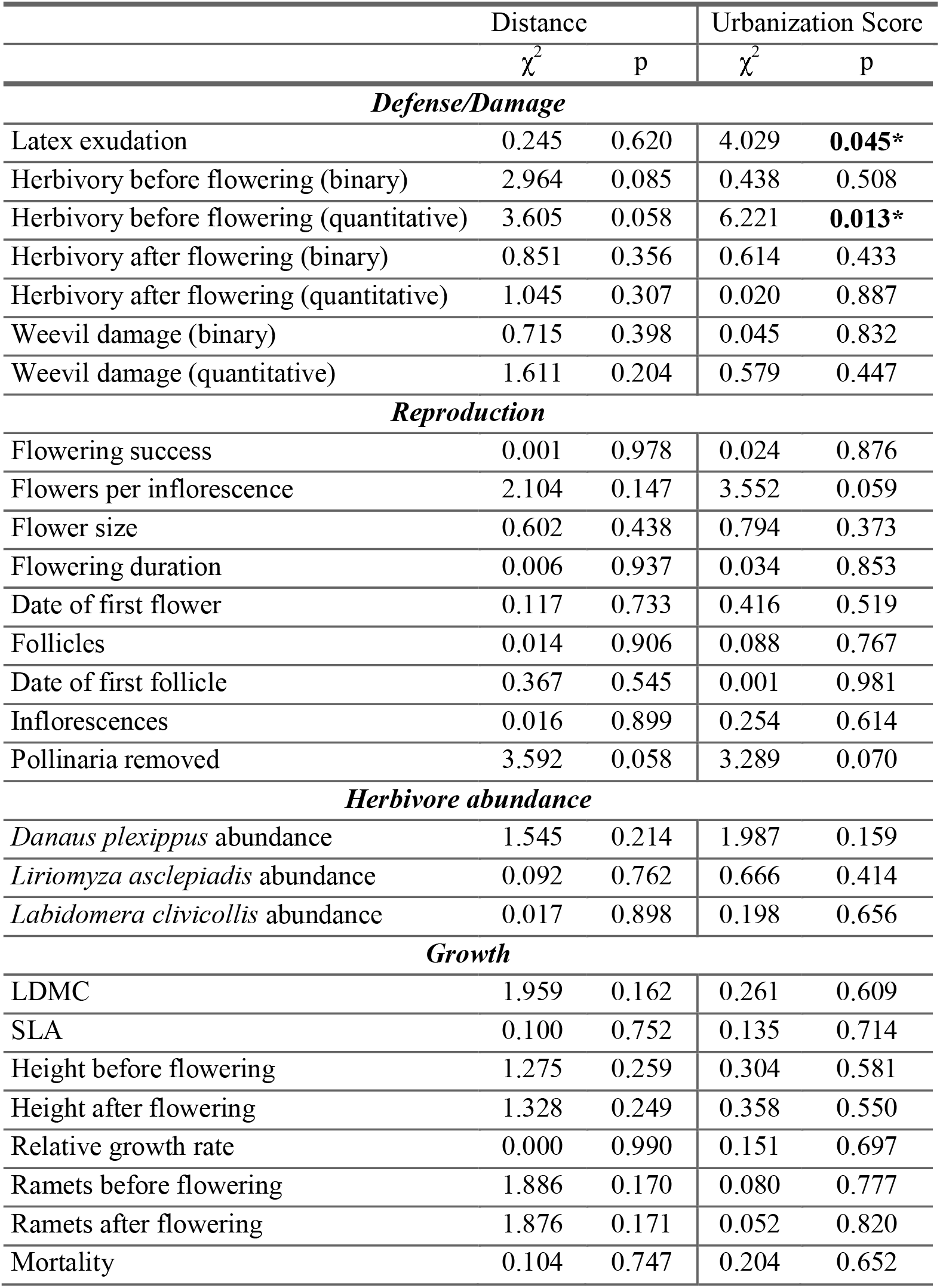
Results from general and generalized linear mixed models examining the effects of urbanization on all phenotypic traits. All populations were included. Shown are maximum likelihood χ^2^ and p-values obtained from type III sums-of-squares ANOVA. Though not shown, block was included as a fixed effect and often explained significant variation in the common garden.

## Results

### Genetic variation within and between populations (Q1)

There was heritable variation for multiple phenotypic traits within populations and evidence of genetic differentiation between populations for some traits. At least one trait per category exhibited significant heritable genetic variation within populations (eight traits, total).

Heritabilities and coefficients of genetic variation ranged from 0-0.588 (mean ± SE: 0.110 ± 0.031) and 0-4.456 (mean ± SE: 0.467 ± 0.207), respectively (Table 1, Supplementary Table S4). The highest statistically significant heritabilities were observed for herbivory after flowering (binary) (*H*² = 0.588, p = 0.004), pollinaria removed (*H*² = 0.544, p = 0.034), and weevil damage (binary) (*H*² = 0.418, p = 0.03), while nine traits, at least one per category, exhibited near-zero heritabilities. Weevil damage (binary) (CV_G_ = 0.995, p = 0.03), herbivory after flowering (binary) (CV_G_ = 0.948, p = 0.004), and *Danaus plexippus* abundance (CV_G_ = 0.767, p = 0.021) exhibited the highest, statistically significant coefficients of genetic variation, while seven traits exhibited near-zero values. It was unlikely that the high frequency of heritable genetic variation within populations was due to chance (binomial expansion test: p < 0.001). Three traits associated with plant defense and reproduction exhibited statistically significant genetic divergence among populations: latex exudation (Q_ST_ = 0.174, p = 0.016), flowering success (i.e., whether plants flowered) (Q_ST_ = 0.492, p = 0.014), and no. of inflorescences (Q_ST_ = 1, p = 0.035) (Table 1, Supplementary Table S4). Q_ST_ ranged from 0 to 1 (mean ± SE: 0.241 ± 0.068) with seven traits exhibiting near-zero values. The relatively few instances of genetic divergence between populations could be due to chance (binomial expansion test: p = 0.15). Thus, these results suggest moderate heritable genetic variation within populations and at most weak phenotypic divergence among populations, both of which are prerequisites for adaptation along an environmental cline.

### Genetic clines along an urban-rural gradient (Q2)

We found little evidence that urbanization influenced genetic divergence between populations. When quantified as distance to the city center, urbanization did not significantly impact phenotypic traits (Fig. 2, Supplementary Figures S4-S12, Table 2, Supplementary Tables S6-S8). When quantified as urbanization score, we detected relationships with latex exudation ( ^2^ = 4.029, p = 0.045, R^2^_m_ = 0.037) and herbivory before flowering (quantitative) (χ^2^ = 6.221, p = 0.013, R^2^_m_ = 0.01). When adjusted for false discovery rates, these positive results were likely due to chance and thus unlikely to be biologically real. Moreover, the multivariate analysis indicated that urbanization did not impact overall phenotype when urbanization was quantified as distance to the city center (F_1,40_ = 14.771, p = 0.711) or urbanization score (F_1,40_ = 6.006, p = 0.984).

**Figure 2.**
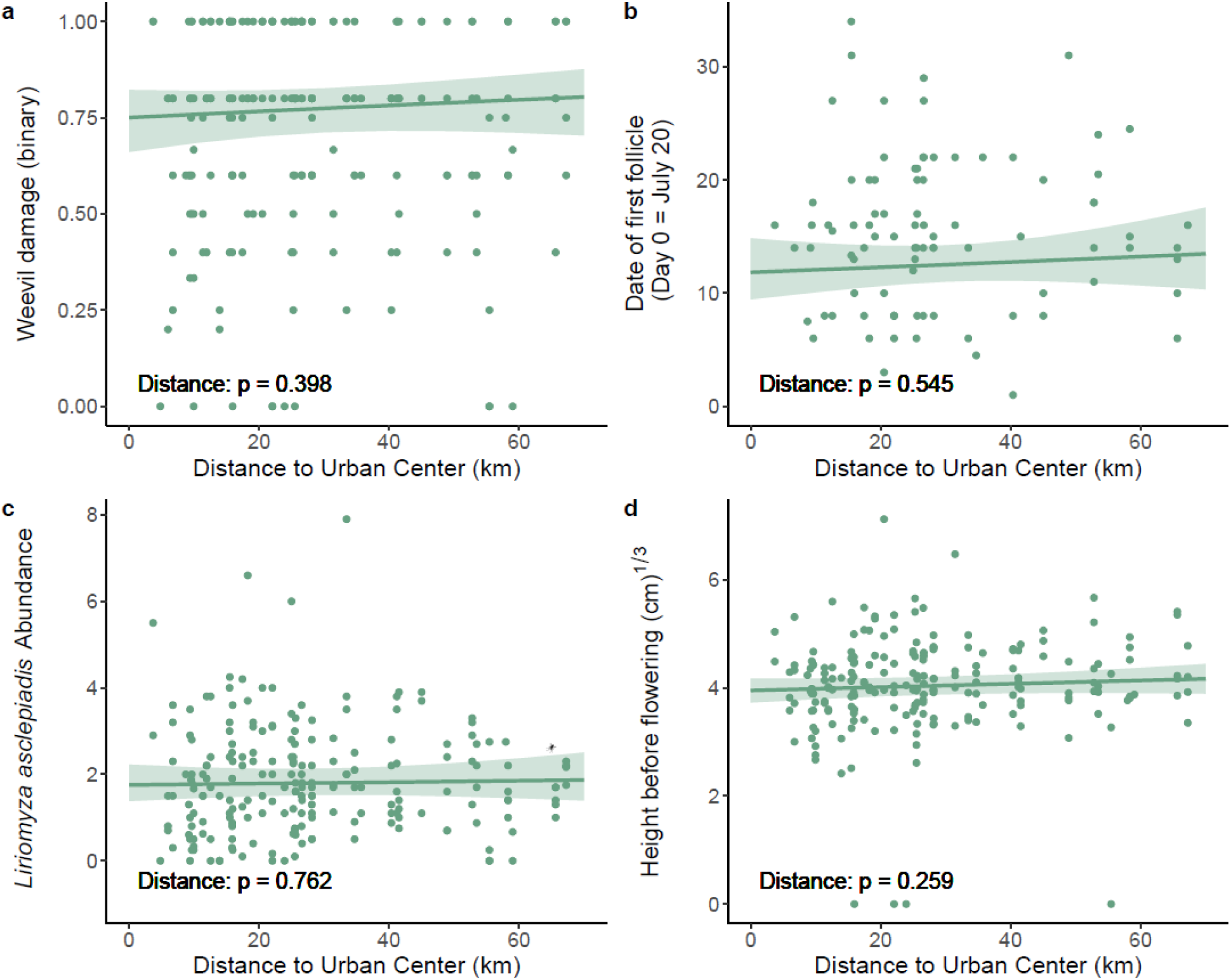
The effect of urbanization on representative traits from each main category: a) plant defense/damage (e.g., the presence of *Rhyssomatus lineaticollis* damage); b) plant reproduction (e.g., date of first follicle); c) herbivore abundance (e.g., *Liriomyza asclepiadis* abundance); and d) plant growth (e.g., height before flowering) when urbanization was quantified by distance from the urban center and all populations were included. Regression lines represent predicted values with a 95% confidence interval and points which represent family-level means are shown for general and generalized linear mixed effects models. These traits were modelled using the following distributions: binomial (a), negative binomial (b, c), and Gaussian (d).

Consistent with our results that urbanization had little effect on genetic divergence, the variation explained and statistical significance of the effects of population and genetic family did not substantially change once urban metrics were included in models (Supplementary Table S5). Taken together, these multiple lines of testing reveal little support for genetic divergence along an urbanization gradient in *A. syriaca*.

### Genetic differentiation between a green corridor & urban matrix (Q3)

Proximity to a green corridor did not strongly influence genetic divergence in phenotypic traits among populations, or the presence of clines along an urbanization gradient (Fig. 3, Supplementary Figures S13-S21, Table 3, Supplementary Tables S7 & S10). Proximity to a green corridor impacted flowers per inflorescence when urbanization was quantified as distance to the city center (χ^2^ = 4.246, p = 0.039, R^2^_m_ <0.001) and urbanization score (χ^2^ = 5.572, p = 0.018, R^2^_m_ <0.001). We detected an interaction between urbanization (quantified as urbanization score) and proximity to a green corridor for latex exudation (χ^2^ = 5.164, p = 0.023, R^2^_m_ = 0.06), flowering duration (χ^2^ = 7.614, p = 0.006, R^2^_m_ = 0.003), follicles (χ^2^ = 7.454, p = 0.006, R^2^ = 0.325), date of first follicle (χ^2^ = 4.467, p = 0.035, R^2m^ = 0.167), inflorescences (χ^2^ = 5.252, p = 0.022, R^2^_m_ <0.001), and weevil damage (binary) (χ^2^ = 4.667, p = 0.031, R^2^ = 0.034). These interactions suggest that the impact of corridors depends on the intensity of urbanization. When adjusted for false discovery rates, these positive results were likely due to chance and the multivariate analysis indicated that proximity to a green corridor did not impact overall phenotypic divergence when urbanization was quantified as distance to the city center (F_1,27_ = 0.346, p = 1) or urbanization score (F_1,27_ = 0.756, p = 1). Consistent with our results that proximity to a green corridor had little effect on genetic divergence, the variation explained and significance of the effects of population and genetic family did not substantially change once urban metrics and proximity to a green corridor were included in models (Supplementary Table S9). These results also reveal little support that proximity to a green corridor is associated with genetic divergence among urban *A. syriaca* populations.

**Table 3.** Results from general and generalized linear mixed models examining the effects of urbanization and proximity to the urban green corridor on all phenotypic traits. Urbanization was quantified via distance from the urban center and only urban populations were included. Shown are maximum likelihood χ^2^ and p-values obtained from type III sums-of-squares ANOVA. Though not shown, block was included as a fixed effect and often explained significant variation in the common garden.

|  | Distance |  | Subtransect |  | D x S |  |
| --- | --- | --- | --- | --- | --- | --- |
| | $\chi^2$ | p | $\chi^2$ | p | $\chi^2$ | p |
| ***Defense/Damage*** |  |  |  |  |  |  |
| Latex exudation | 1.612 | 0.204 | 0.014 | 0.906 |  |  |
| Herbivory before flowering (binary) | 1.023 | 0.312 | 0.462 | 0.497 |  |  |
| Herbivory before flowering (quantitative) | 0.054 | 0.817 | 0.122 | 0.727 |  |  |
| Herbivory after flowering (binary) | 0.581 | 0.446 | 0.672 | 0.412 |  |  |
| Herbivory after flowering (quantitative) | 0.004 | 0.949 | 0.389 | 0.533 |  |  |
| Weevil damage (binary) | 3.572 | 0.059 | 0.166 | 0.684 |  |  |
| Weevil damage (quantitative) | 0.082 | 0.774 | 0.829 | 0.363 |  |  |
| ***Reproduction*** |  |  |  |  |  |  |
| Flowering success | 0.095 | 0.757 | 0.625 | 0.429 |  |  |
| Flowers per inflorescence | 0.546 | 0.460 | 4.246 | **0.039\*** |  |  |
| Flower size | 1.374 | 0.241 | 0.576 | 0.448 |  |  |
| Flowering duration | 0.410 | 0.522 | 2.027 | 0.155 |  |  |
| Date of first flower | 0.725 | 0.394 | 0.006 | 0.939 |  |  |
| Follicles | 0.003 | 0.953 | 0.580 | 0.446 | 0.098 | 0.755 |
| Date of first follicle | 3.306 | 0.069 | 0.944 | 0.331 |  |  |
| Inflorescences | 0.199 | 0.656 | 0.454 | 0.501 |  |  |
| Pollinaria removed | 0.379 | 0.538 | 0.012 | 0.912 |  |  |
| ***Herbivore abundance*** |  |  |  |  |  |  |
| *Danaus plexippus* abundance | 4.697 | **0.030\*** | 0.158 | 0.691 |  |  |
| *Liriomyza asclepiadis* abundance | 0.290 | 0.590 | 0.168 | 0.681 |  |  |
| *Labidomera clivicollis* abundance | 0.404 | 0.525 | 0.068 | 0.794 |  |  |
| ***Growth*** |  |  |  |  |  |  |
| LDMC | 2.710 | 0.100 | 1.456 | 0.228 |  |  |
| SLA | 0.400 | 0.527 | 0.840 | 0.360 |  |  |
| Height before flowering | 0.126 | 0.722 | 0.009 | 0.925 |  |  |
| Height after flowering | 3.484 | 0.062 | 0.056 | 0.813 | 2.366 | 0.124 |
| Relative growth rate | 0.036 | 0.849 | 0.564 | 0.453 |  |  |
| Ramets before flowering | 2.861 | 0.091 | 0.002 | 0.962 | 2.050 | 0.152 |
| Ramets after flowering | 6.005 | **0.014\*** | 3.231 | 0.072 | 2.740 | 0.098 |
| Mortality | 1.089 | 0.297 | 0.062 | 0.803 |  |  |

**Figure 3.**
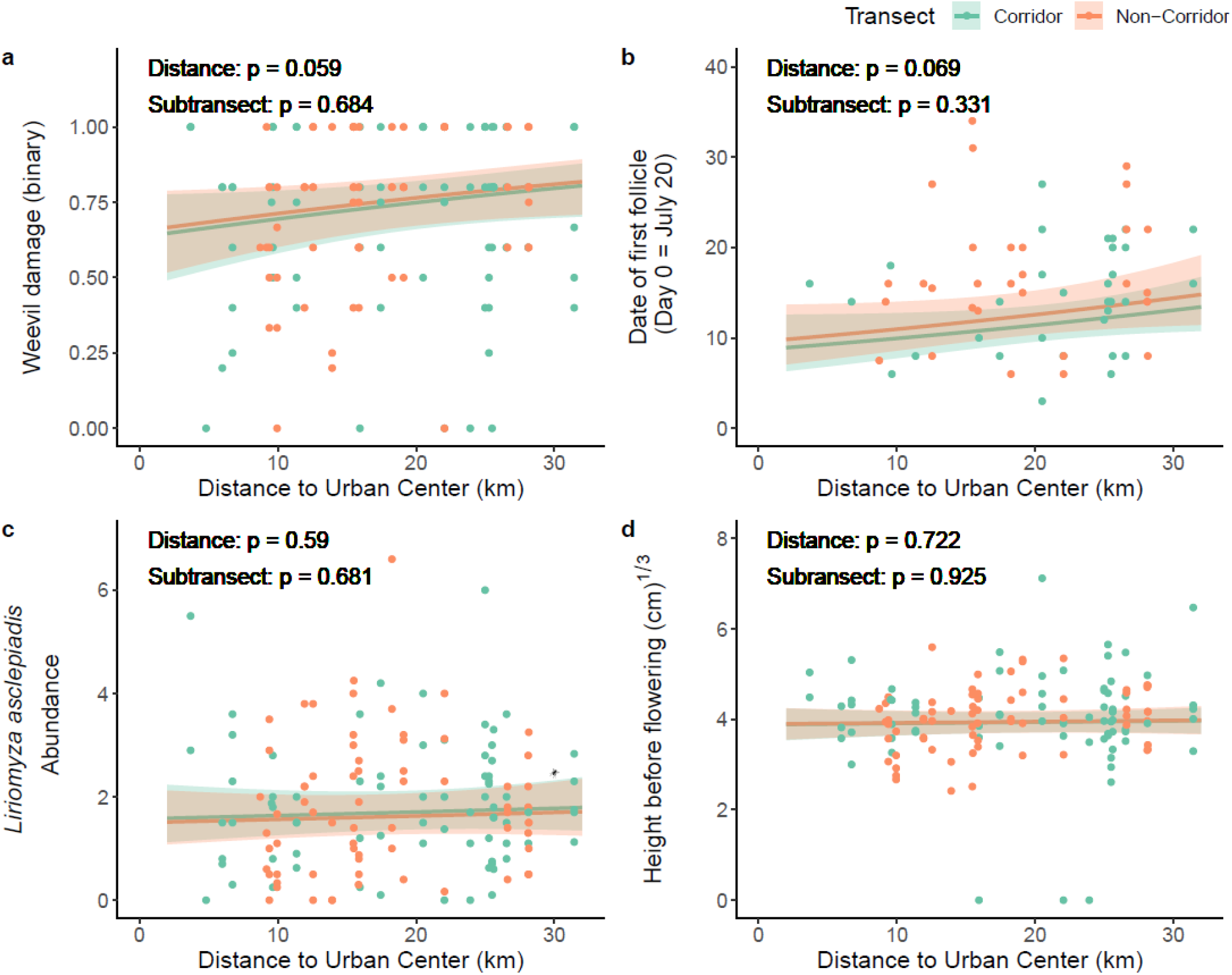
The effects of urbanization and proximity to a green corridor on representative traits from each main category: a) plant defense/damage (e.g., the presence of *Rhyssomatus lineaticollis* damage); b) plant reproduction (e.g., date of first follicle); c) herbivore abundance (e.g., *Liriomyza asclepiadis* abundance); and d) plant growth (e.g., height before flowering) when urbanization was quantified by distance from the urban center and only urban populations were included. Regression lines represent predicted values with a 95% confidence interval and points which represent family-level means are shown for general and generalized linear mixed effects models. These traits were modelled using the following distributions: binomial (a), negative binomial (b, c), and Gaussian (d).

## Discussion

In this study, we tested the hypotheses that urbanization and an urban green corridor drive genetic divergence in phenotypic traits among populations of *A. syriaca*. Though we observed moderate heritable genetic variation within populations and some weak phenotypic divergence among populations, we found low support that urbanization or proximity to an urban green corridor influenced genetic divergence. These results suggest that *A. syriaca* has not undergone rapid evolutionary change in response to urban landscapes as a whole, nor a common component of urban environments: a green corridor. Below, we discuss the implications of these results for understanding how species evolve in response to rapid environmental change in heterogeneous urban environments.

### Genetic variation within and between populations (Q1)

Genetic divergence of phenotypic traits along an urbanization gradient necessitates that populations exhibit genetic divergence, which is more likely to occur when those populations contain heritable phenotypic variation. These criteria have been observed in previous studies of *A. syriaca*, although not in an urban context. For instance, many phenotypic traits are heritable, especially those associated with defense/damage and growth; examples include *Tetraopes spp.* damage, carbon:nitrogen ratio, SLA, water percentage, trichome density, ant abundance, root and shoot mass, constitutive and induced latex and cardenolides^62–66, 89^. In addition, substantial heritable phenotypic variation within populations was found at both a continental scale (i.e., within populations sampled across the species’ native North American range^90^) and a local scale (i.e., within single genetic populations^63, 64^, ^but^ ^see^ ^91^). Similarly, genetic divergence for growth-related traits was detected among *A. syriaca* populations sampled from a latitudinal gradient from New Brunswick, Canada, to North Carolina, USA^66^. In our common garden experiment, we found moderate heritable genetic variation within populations and weak phenotypic divergence among populations sampled from an urbanization gradient. Our results, which indicate that these populations have met the prerequisites to evolve in response to urbanization and/or an urban green corridor, are mostly consistent with previous findings from non-urban contexts.

Our estimates of genetic variation for phenotypic traits within populations are comparable to previous studies. For example, heritability estimates for plant height after flowering and the number of ramets (range: *H*² = 0.132-0.161) were close to estimates by Vannette et al.^65^ (aboveground biomass: *H*² = 0.11) and Agrawal et al.^62^ (vegetative biomass: *H*² = 0.12). We found significant and relatively high heritability estimates for herbivory after flowering (*H*² = 0.588) and weevil damage (*H*² = 0.418) when measured on a presence/absence basis, but not quantitatively when measured as percent leaf area removed. In comparison, Agrawal et al.^62^ found moderate heritability for herbivory when measured as the percentage of leaves with foliar damage due to chewing herbivores (*H*² = 0.284), but not for weevil damage when measured as the length of stem scars (*H*² = 0.037). By calculating quantitative genetic parameter estimates for a diverse group of traits, we estimated heritabilities for traits associated with sexual reproduction— most for the first time in this species. Moreover, we found that two of these traits had moderate to high heritabilities (i.e., pollinaria removed: *H*² = 0.544; date of first follicle: *H*² = 0.235), suggesting a higher propensity for these traits to evolve in response to environmental change. Overall, these results confirm the capacity for these *A. syriaca* populations to evolve in response to environmental stressors, such as urbanization.

### Genetic clines along an urban-rural gradient (Q2)

Many taxa exhibit genetic divergence between urban and nonurban populations for various traits. In plants, this has been documented for traits associated with phenology^16, 27, 29, 30^, size^16, 29^, fecundity^16, 29^, defense^16^, and competitive ability^92^. Additionally, in both native and introduced ranges, *A. syriaca* exhibits clines for growth and leaf physiology traits that correspond with a defining feature of urban environments: temperature^90^. Yet despite surveying several suites of traits in *A. syriaca*, we found low support for genetic divergence along an urbanization gradient. Furthermore, we detected only small effect sizes (range: 0.01-0.037) for the few traits associated with urbanization even with the large scale and replication afforded by our experimental design. Thus, multiple lines of evidence suggest the lack of such divergence at present, though we do not rule out the possibility of genetic divergence emerging in the future.

The relative rarity of this outcome in the existing urban evolution literature presents a valuable opportunity to explore circumstances that could prevent urbanization from influencing genetic divergence among populations. In this case, evolutionary change may have been precluded by urban ecological pressures that were possibly too small or brief, as the city of Toronto has only had a population ≥50,000 residents for ca. 150 years^93^. The life history traits of *A. syriaca* could also slow evolutionary change. For example, the vegetative reproduction inherent to *A. syriaca* can lead to clonal growth and the loss of genotypic diversity within populations^94, 95^, which is compounded by the species’ long-lived nature. Also, given that *A. syriaca* is largely self-incompatible, wind-dispersed, and frequently pollinated by insects that can travel >1 km (e.g., *Bombus* spp*., Apis mellifera*)^96, 97^, long-distance pollen and seed flow could yield high gene flow among populations and low genetic drift within populations. Lastly, additional anthropogenic factors such as noxious weed legislation^98^ may have depleted genetic diversity in natural populations, while imported nursery stock could have introduced nonlocal alleles into the gene pool, potentially slowing the rate of adaptation.

This study lends valuable context to prior work demonstrating how urbanization impacts the reproductive success of *A. syriaca* populations^48^. Specifically, as we observed relatively little phenotypic divergence among populations within the common garden, our results suggest that the previously observed *in-situ* phenotypic divergence was mostly consistent with phenotypic plasticity as opposed to genetic divergence. This finding underscores the importance of investigating the genetic basis of phenotypic divergence observed in cities and reiterates the role of phenotypic plasticity in shaping how populations respond to anthropogenic stress. Future studies could investigate the specific conditions that promote phenotypic plasticity within urban environments.

### Genetic differentiation between a green corridor & urban matrix (Q3)

Very little is known about how urban green corridors influence genetic divergence among plant populations. In non-urban environments, green corridors are predicted to increase gene flow among plant populations by facilitating pollen flow and seed dispersal, which is expected to decrease genetic divergence between populations^99–101^. Limited research in urban environments suggests that green corridors often increase gene flow among animal populations^46, 102–104^, ^but^ ^see^ ^105^, and that urban features such as railways can function as corridors among plant and animal populations^106, 107^. In our study, proximity to a green corridor did not strongly influence genetic divergence in phenotypic traits among *A. syriaca* populations.

As discussed above for Question 2, urbanization may not restrict gene flow among populations. If true, there would be little opportunity for the green corridor to significantly enhance gene flow and diminish any hypothetical genetic divergence in *A. syriaca*, as proposed for butterflies in Angold et al.^105^. Conversely, our data also suggest that proximity to a green corridor does not inherently facilitate genetic divergence either. It is also possible that urban green corridors actually impact genetic divergence but that our results reflect the specific environmental conditions associated with our chosen corridor; i.e., the efficacy of the corridor could have been impeded by a highly hostile external matrix or edge effects generated by the narrow shape^108, 109^. Relatedly, the corridor we sampled would have provided minimal connectivity to plants or pollen vectors if it were not perceived as a functional corridor to either group. Regardless, these findings contradict the expectation that habitat fragmentation impacts the evolutionary processes of urban populations and invite further research into how and when corridors impact genetic divergence of phenotypic traits in urban populations.

## Conclusion

Urbanization is associated with phenotypic trait divergence for many species, yet in most cases, the genetic basis of these trait differences is unresolved. Additionally, the extent to which specific aspects of the urban landscape impact genetic divergence in phenotypic traits among plant populations is virtually unknown. Here, we show that neither urbanization nor an urban green corridor impacted genetic divergence in phenotypic traits in common milkweed, a native plant of conservation importance. These results demonstrate an example in which urbanization has not substantially contributed to genetic divergence among populations. To our knowledge, this study is also the first to investigate if urban green corridors impact genetic divergence in phenotypic traits in plant populations and ultimately suggests the absence of such effects in our study system. To further understand how urban environments and urban green corridors impact eco-evolutionary dynamics in plant populations, future research should verify the consistency of these findings by testing similar questions in plants with diverse life history, and in a range of cities with heterogeneous landscapes.

## Acknowledgements

We thank the following people who helped maintain the experiment and collect data: L. Albano, K. Brown, B. Cohan, S. El-Galmady, H. Fargo, I. Fortin, S. Innes, M. Irving, A. Joyce, D. Karimov, S. Kharab, K. Manikonda, L. Miles, R. Molnarova, D. Murray-Stoker, N. Naik, N. Navaratne, V. Nhan, M. Oman, V. Quiroga Angel, R. Rivkin, C. Roux, J. Santangelo, J. Stinchcombe, W. Sturch, T. Tejal, A. Tomchyshyn, O. Toth, and Y. Yu. A. Hastings performed the cardenolide analysis, and L. Albano, A. Caizergues, and D. Murray-Stoker provided feedback on manuscript drafts. This work was funded by a University of Toronto EEB PhD Grant (STB), University of Toronto Mississauga Centre for Urban Environments Research Grant (STB), a Grant-In-Aid of Research from Sigma Xi, The Scientific Research Society (STB), the NSERC CREATE program “ADVENT/ENVIRO” (Murray et al.) (STB), NSERC Discovery Grants (HHW & MTJJ), a Canada Research Chair (MTJJ), and an E.W.R. Steacie Fellowship (MTJJ). We also wish to acknowledge this land on which the University of Toronto operates. For thousands of years it has been the traditional land of the Huron-Wendat, the Seneca, and the Mississaugas of the Credit. Today, this meeting place is still the home to many Indigenous people from across Turtle Island and we are grateful to have the opportunity to work on this land. It is a privilege for us to perform research on a plant which has traditionally been a source of food, medicine, and fiber for many Indigenous communities, including those who have been caretakers of this land for time immemorial.

## Author contributions

STB, MTJJ, and HHW conceived the project. All authors designed the study. STB and MTJJ conducted the field work. AAA collected the cardenolide data. STB analyzed the data, wrote the manuscript, and revised it with input from AAA, MTJJ, and HHW.

## Data availability statement

Data are archived on Zenodo (DOI: 10.5281/zenodo.7615892). Code will be submitted upon acceptance of the manuscript.

