## Supplemental File for "Urbanization and a green corridor do not impact genetic divergence in common milkweed (*Asclepias syriaca*)"

Contents:

- Supplementary text S1
- Supplementary tables and figures:
    - Supplementary figures: Figures S1-S21
    - Supplementary tables: Tables S1-S10

**Text S1**

***Common garden experiment***

Full-sibling seeds were cleaned with 5% bleach, mechanically scarified with a razor, vernalized via refrigeration at 4°C for a week, and incubated at 30°C for 3-4 days until germination occurred. Upon germination, two full-sib seeds were planted in the same pot with potting soil (Pro-Mix LP15, Sun Gro Horticulture) and Nutricote 14:13:13 (N:P:K) slow release fertilizer (Plant Products Inc., Ancaster, Canada). Pots were placed in a growth chamber set to 27°C daytime/25°C nighttime temperatures with 14h light:10h dark and 750 μmol/m²/s of light with 50% humidity. Plants were watered daily and randomized one week after planting. All pots were thinned to one seedling after two weeks. If both seeds sprouted two weeks after planting, one seedling was wholly removed.

Once brought to KSR, pots were sunk into the ground within a hole cut in the fabric. Seedlings were watered immediately after transplanting as well as once during the first week after transplanting and four weeks after transplanting due to unusually hot and dry conditions. Vegetated laneways next to rows were routinely mowed throughout the growing season (May-September). Pots were weeded annually and otherwise grown under natural conditions for the remainder of the experiment. Landscaping fabric was replaced in 2020 (Hanes Geo Components, Winston-Salem, NC, USA) due to degradation.

***Trait measurements***

We generated population-level estimates of leaf cardenolide concentrations by collecting the leaf used to assess latex exudation and its opposite, placing leaves on dry ice, and storing at -80°C until freeze-dried. We cut tissue from each replicate, excluding the mid-rib, and pooled the samples by population so that each population’s vial contained 50 mg of tissue from 5 replicates per family, and 5 families per population except when mortality prevented collection from all plants within a family or population. For instance, we collected 2 mg/replicate if a population contained the full 25 replicates; otherwise, we collected >2mg/replicate for up to 10 mg/family for populations containing <25 replicates.

To assess specific leaf area (SLA) and leaf dry matter content (LDMC), we collected the youngest fully expanded intact leaf, placed it in a coin envelope on dry ice, and saturated the leaves in water for 13 h. Leaves were dabbed with a paper towel, photographed with a ruler for scale, weighed for wet mass, and then dried at 60°C for 48 hours, after which leaves were re-weighed for dry mass. Leaf length, width, and area were measured with ImageJ[1].

***Statistical analyses***

*Genetic differentiation between a green corridor & urban matrix (Q3)*

For Questions 2 and 3, we initially fitted multi-year models with the addition of Year as a fixed effect. We fitted 1-year models by removing the effect of year from the multi-year models and restricting the data to the last year of sampling, then compared the significance of each fixed effect between analogous multi-year and 1-year models. When we tested for the consistency of urbanization and green corridors across all years of data collection, we found that these effects were qualitatively identical to those reported in Tables 2-3 85% of the time (Supplementary Table 3).

### Figures

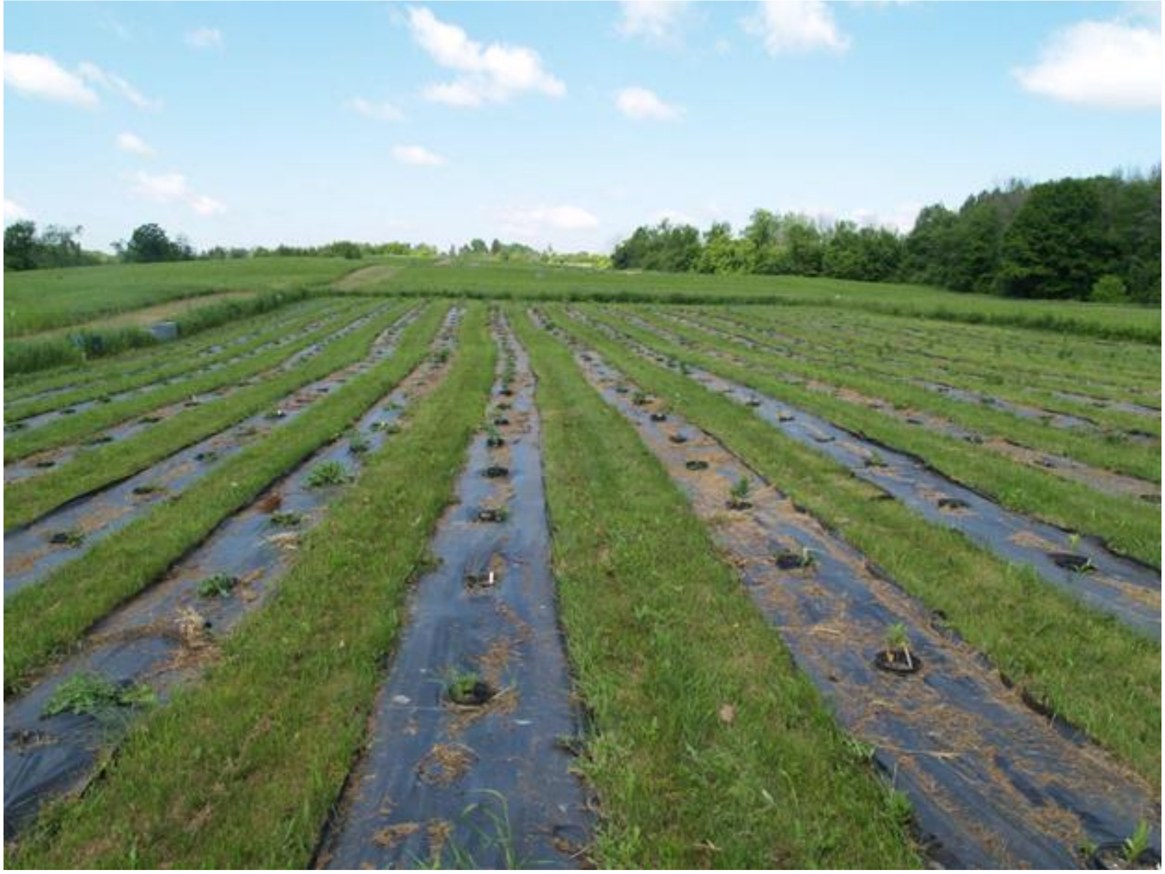

**Supplementary Figure 1.** Common garden experiment setup. Photo credit: Sophie Breitbart

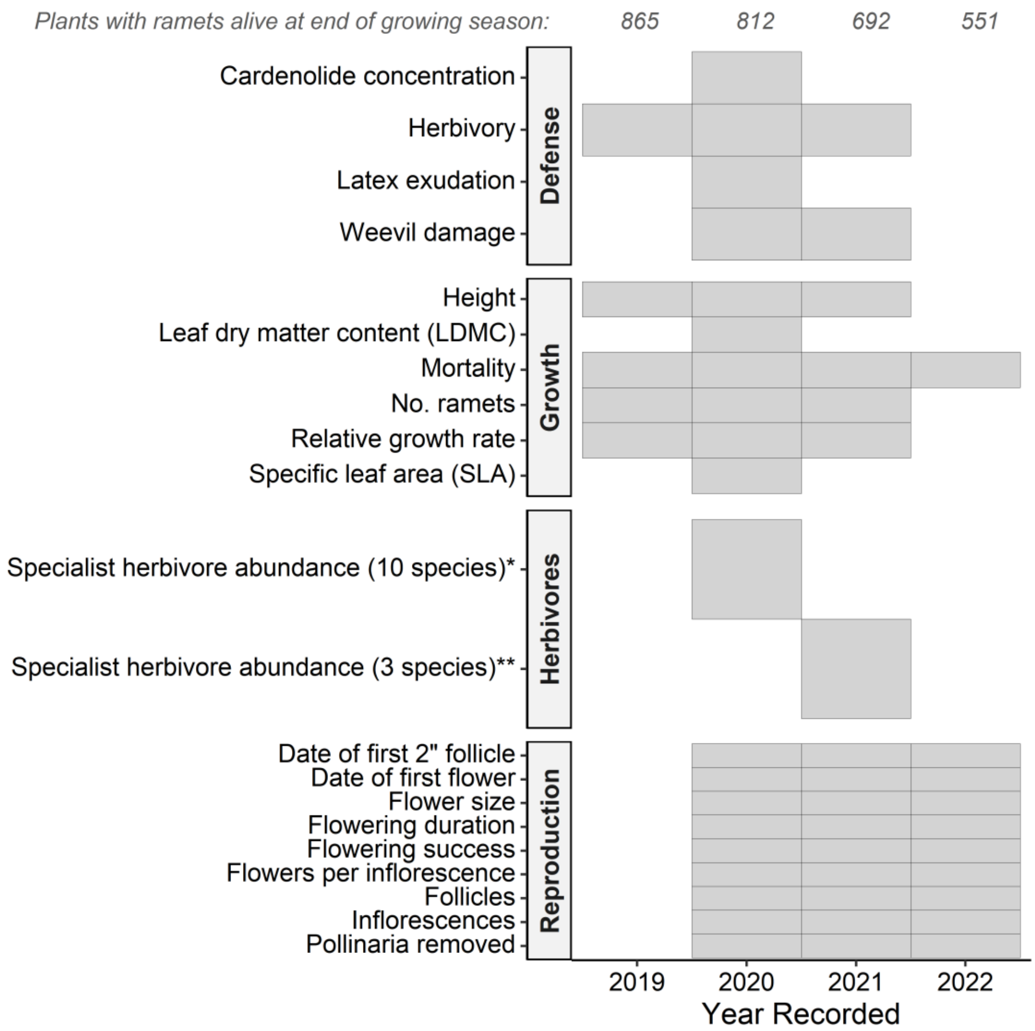


**Aphis asclepiadis, Aphis nerii, Danaus plexippus, Euchaetes egle, Labidomera clivicollis, Liriomyza asclepiadis, Lygaeus kalmii, Myzocallis asclepiadis, Rhyssomatus lineaticollis, Tetraopes tetropthalmus*

***D. plexippus, L. clivicollis, L. asclepiadis*

**Supplementary Figure 2.** Observation schedule for traits associated with plant reproduction, growth, and defense/damage throughout 2019-2022 field seasons and number of plants alive per year.

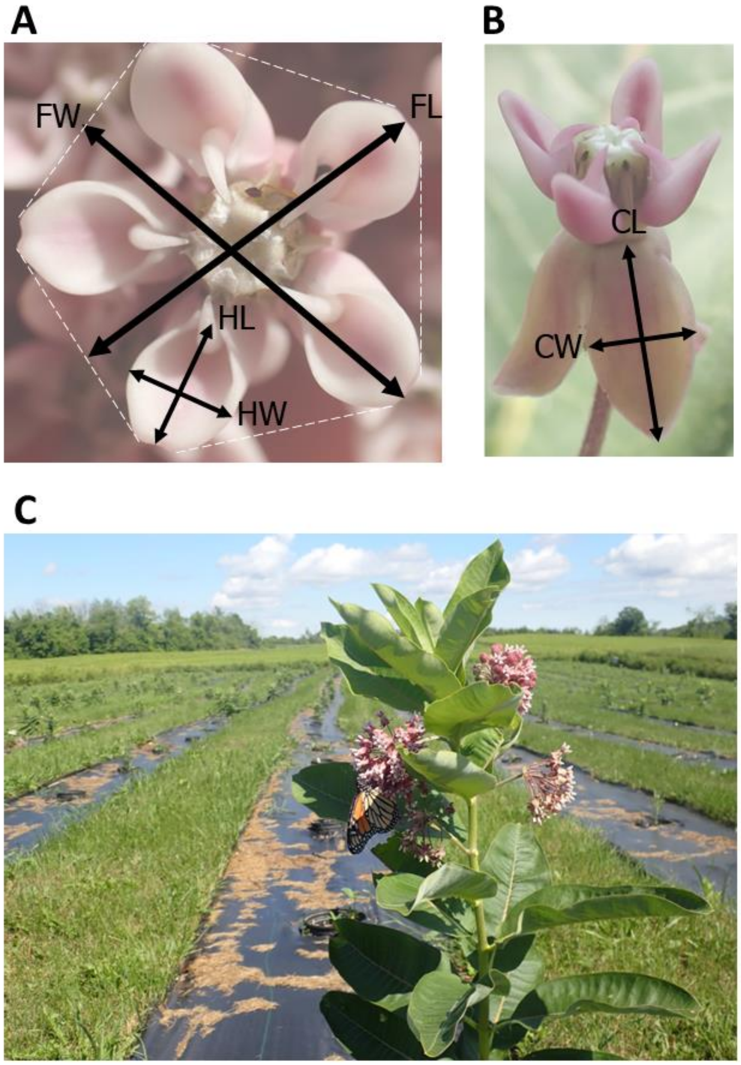


**Supplementary Figure 3.** A) Example of hood length (HL), hood width (HW), flower length (FL), and flower width (FW) measurements. B) Example of corolla length (CL) and corolla width (CW) measurements. All six measurements were averaged to calculate mean size per flower. C) A flowering *A. syriaca* ramet within the common garden. Photo credit: Sophie Breitbart

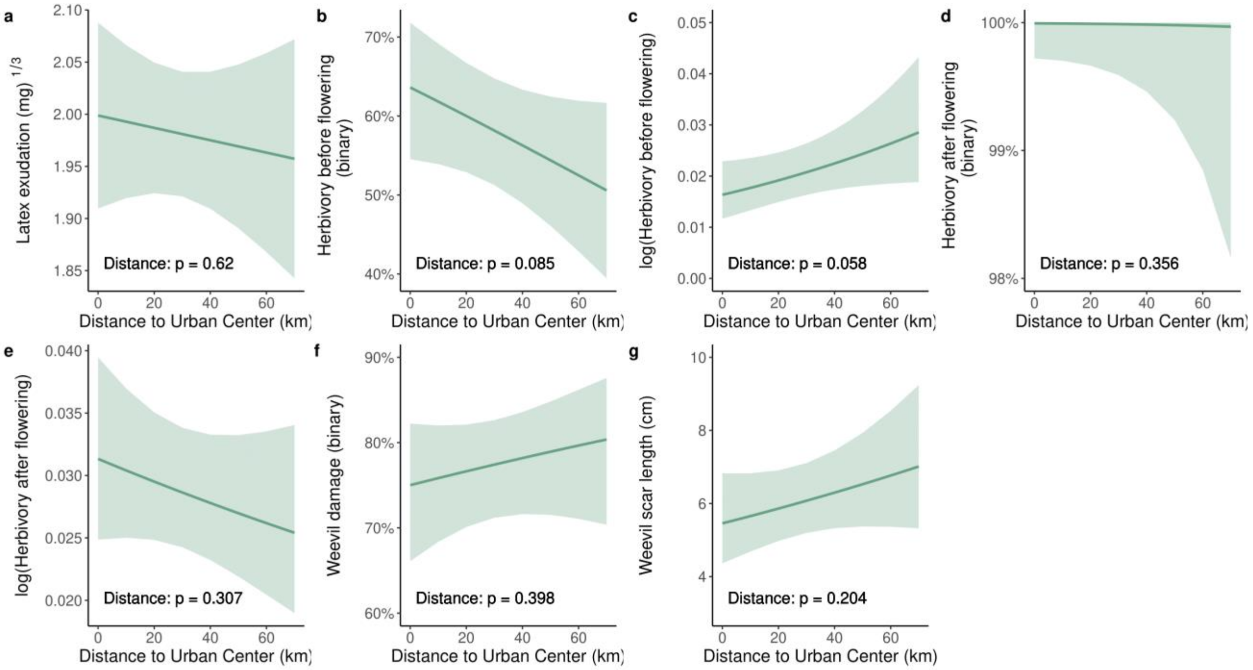


**Supplementary Figure 4.** The effect of urbanization on plant defense/damage traits when urbanization was quantified by distance from the urban center. Regression lines with a 95% confidence envelope for the mean response are shown for general and generalized linear mixed effects models.

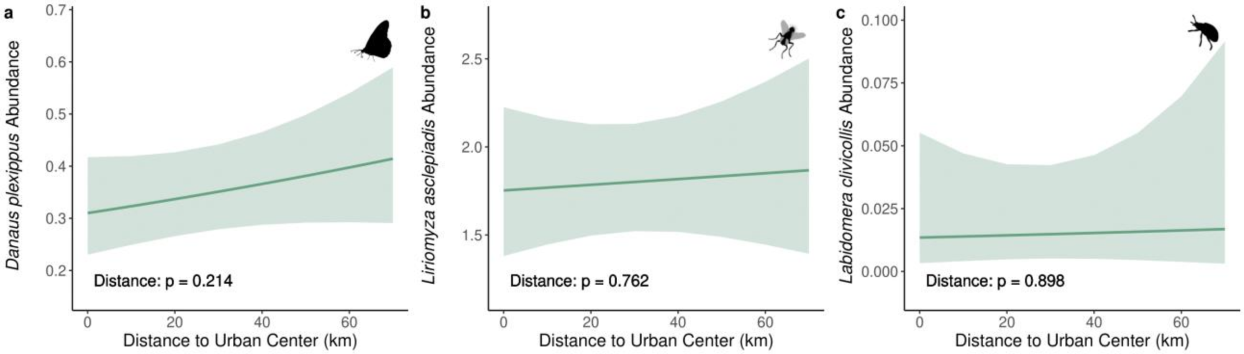


**Supplementary Figure 5.** The effect of urbanization on herbivore abundance when urbanization was quantified by distance from the urban center. Regression lines with a 95% confidence envelope for the mean response are shown for generalized linear mixed effects models.

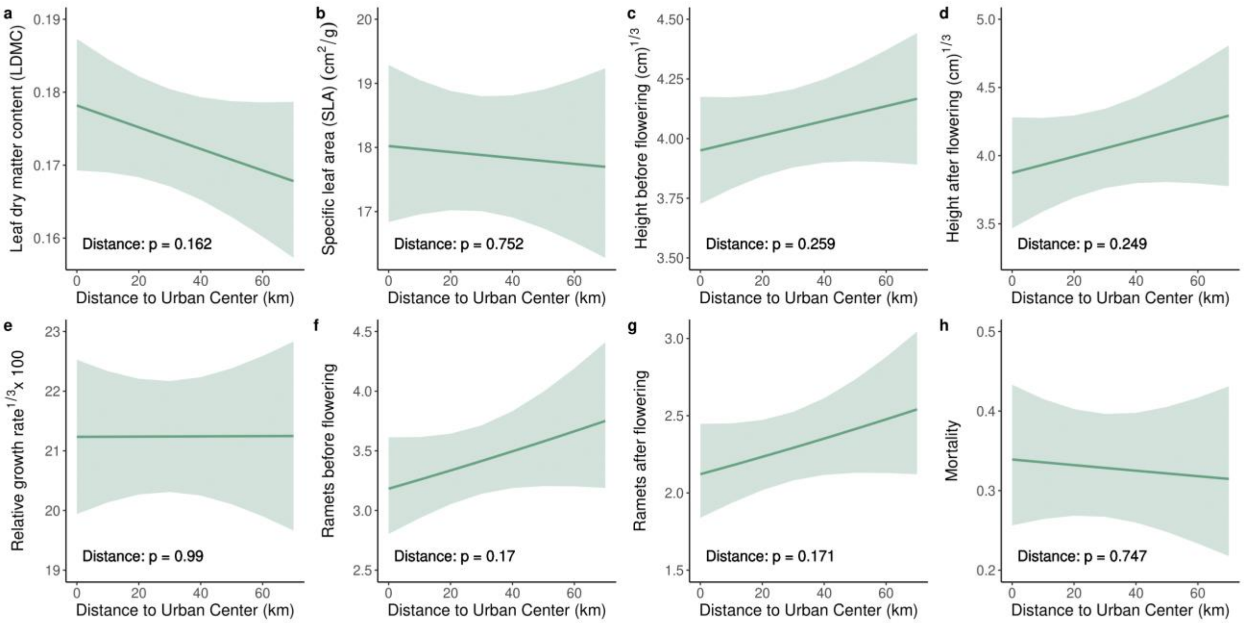


**Supplementary Figure 6.** The effect of urbanization on plant growth traits when urbanization was quantified by distance from the urban center. Regression lines with a 95% confidence envelope for the mean response are shown for general and generalized linear mixed effects models.

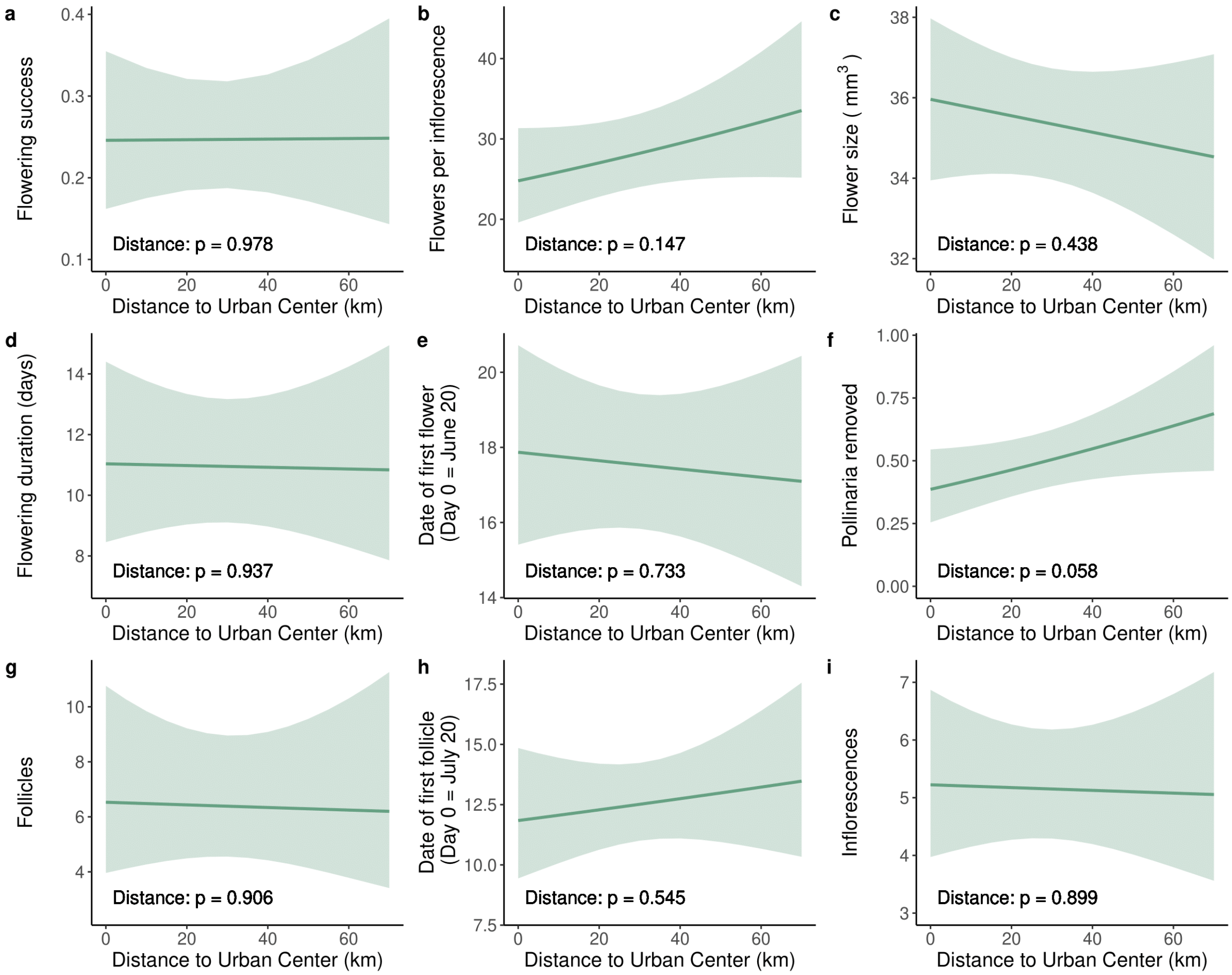


**Supplementary Figure 7.** The effect of urbanization on plant reproduction traits when urbanization was quantified by distance from the urban center. Regression lines with a 95% confidence envelope for the mean response are shown for general and generalized linear mixed effects models.

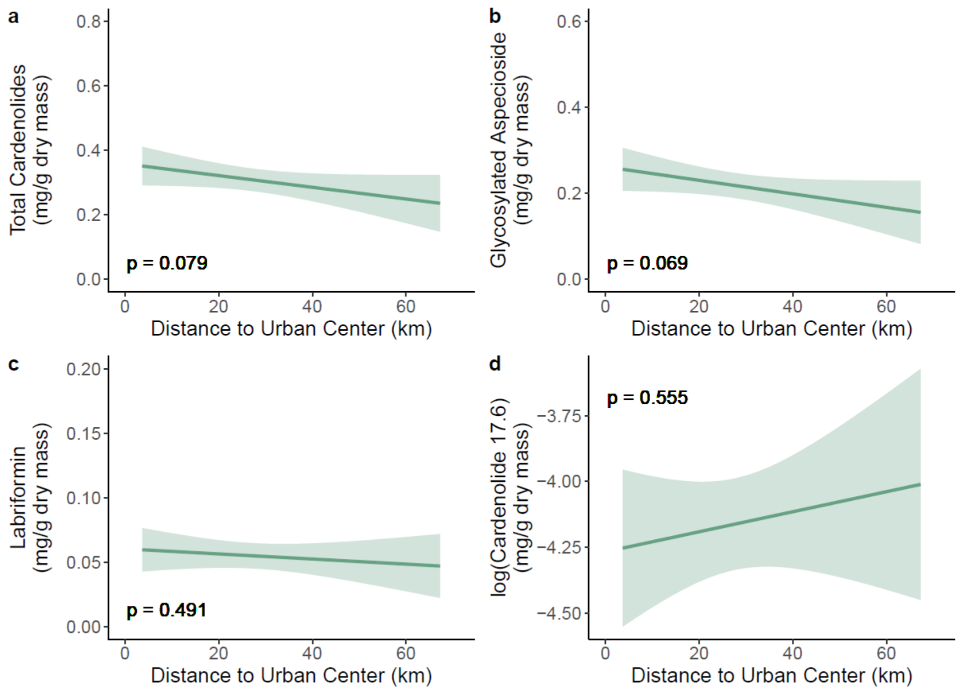


**Supplementary Figure 8.** The effect of urbanization on cardenolides when urbanization was quantified by distance from the urban center. Regression lines with a 95% confidence envelope for the mean response are shown for general linear mixed effects models. Cardenolide 17.6 is an unidentified cardenolide with a retention time of 17.6 minutes.

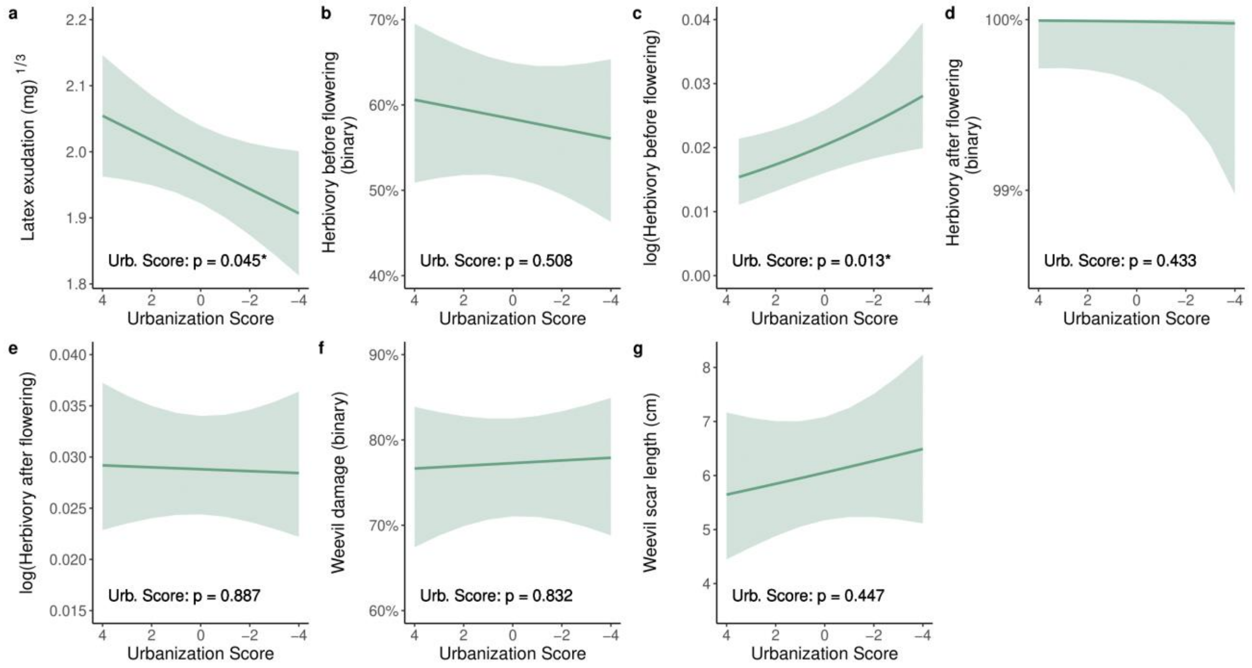


**Supplementary Figure 9.** The effect of urbanization on plant defense/damage traits when urbanization was quantified by urbanization score. Regression lines with a 95% confidence envelope for the mean response are shown for general and generalized linear mixed effects models.

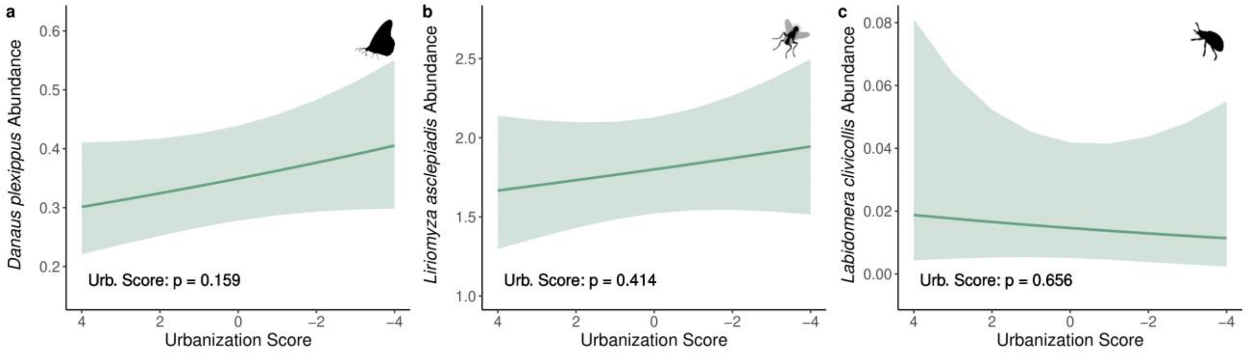


**Supplementary Figure 10.** The effect of urbanization on herbivore abundance when urbanization was quantified by urbanization score. Regression lines with a 95% confidence envelope for the mean response are shown for generalized linear mixed effects models.

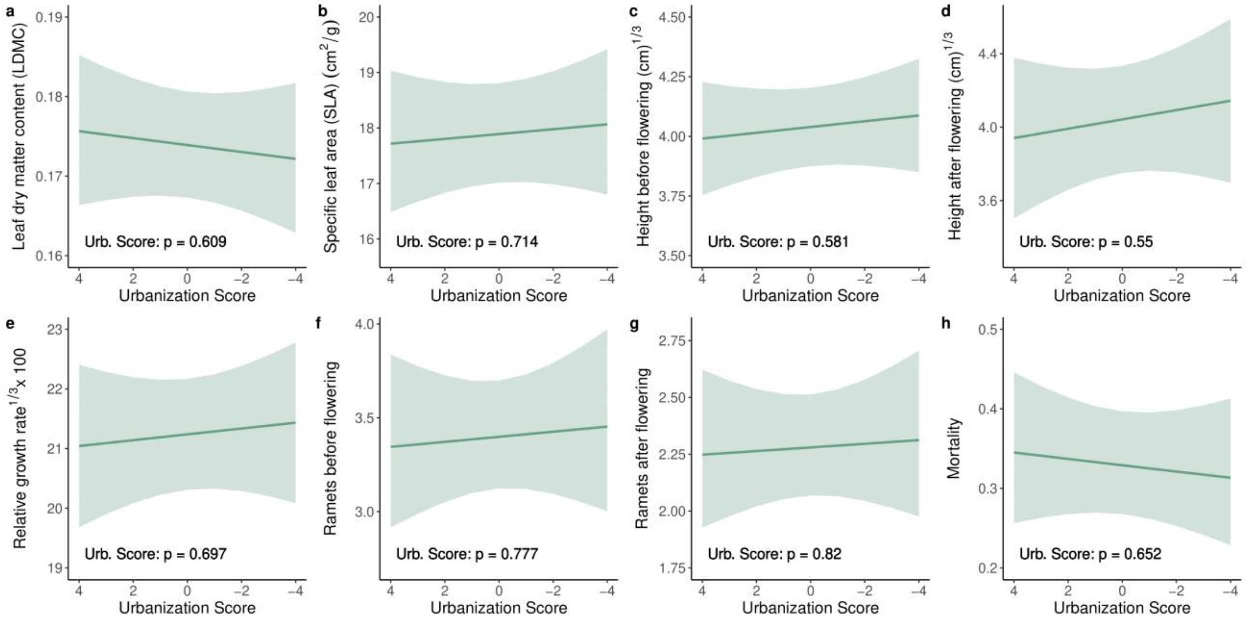


**Supplementary Figure 11.** The effect of urbanization on plant growth traits when urbanization was quantified by urbanization score. Regression lines with a 95% confidence envelope for the mean response are shown for general and generalized linear mixed effects models.

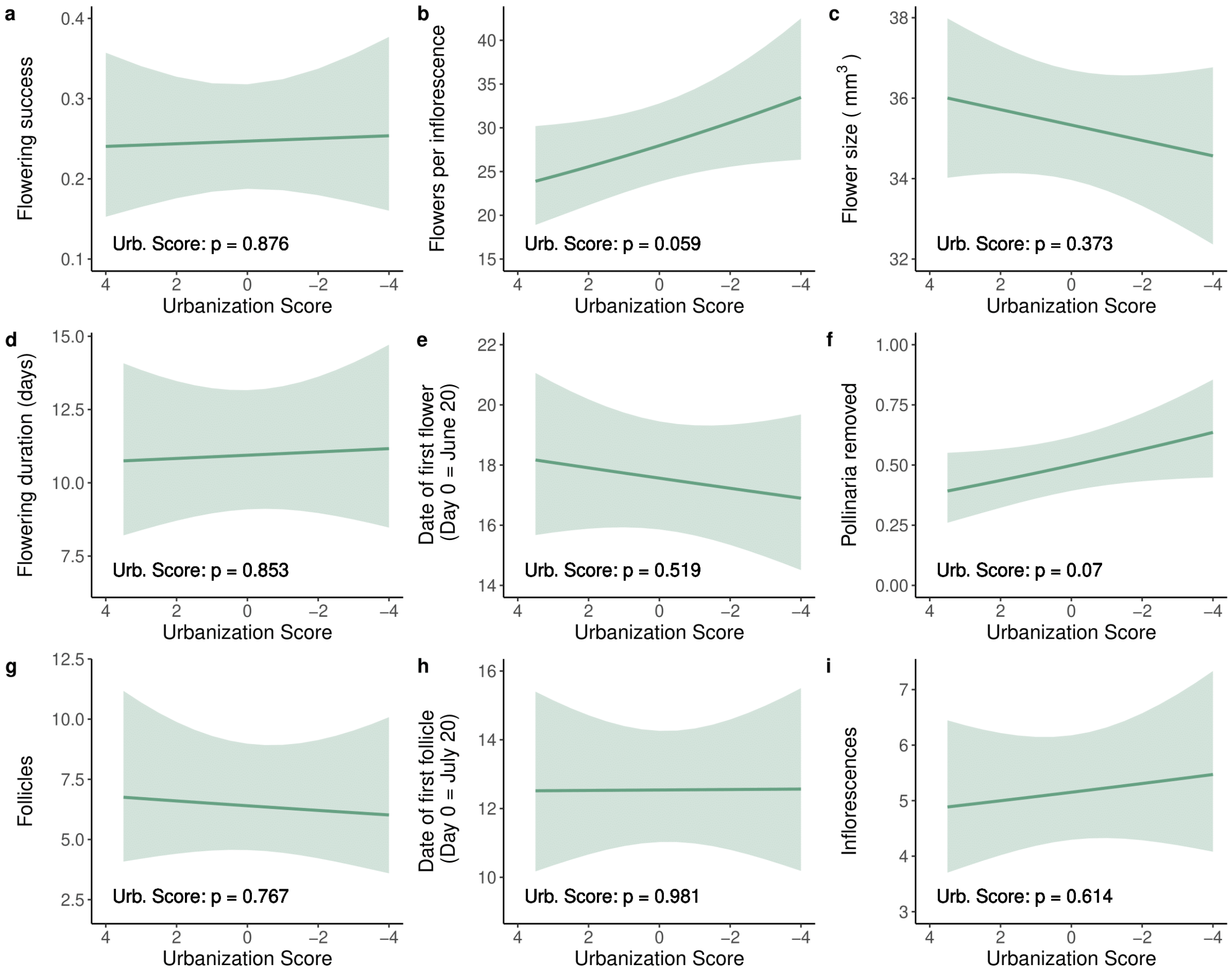


**Supplementary Figure 12.** The effect of urbanization on plant reproduction traits when urbanization was quantified by urbanization score. Regression lines with a 95% confidence envelope for the mean response are shown for general and generalized linear mixed effects models.

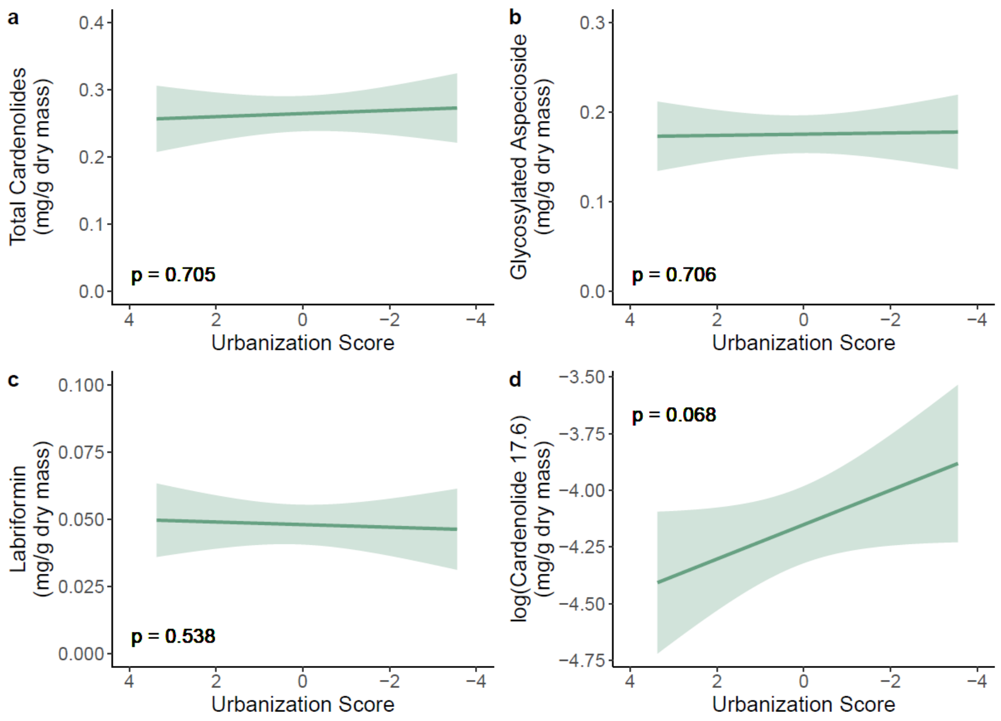


**Supplementary Figure 13.** The effect of urbanization on cardenolides when urbanization was quantified by urbanization score. Regression lines with a 95% confidence envelope for the mean response are shown for general linear mixed effects models. Cardenolide 17.6 is an unidentified cardenolide with a retention time of 17.6 minutes.

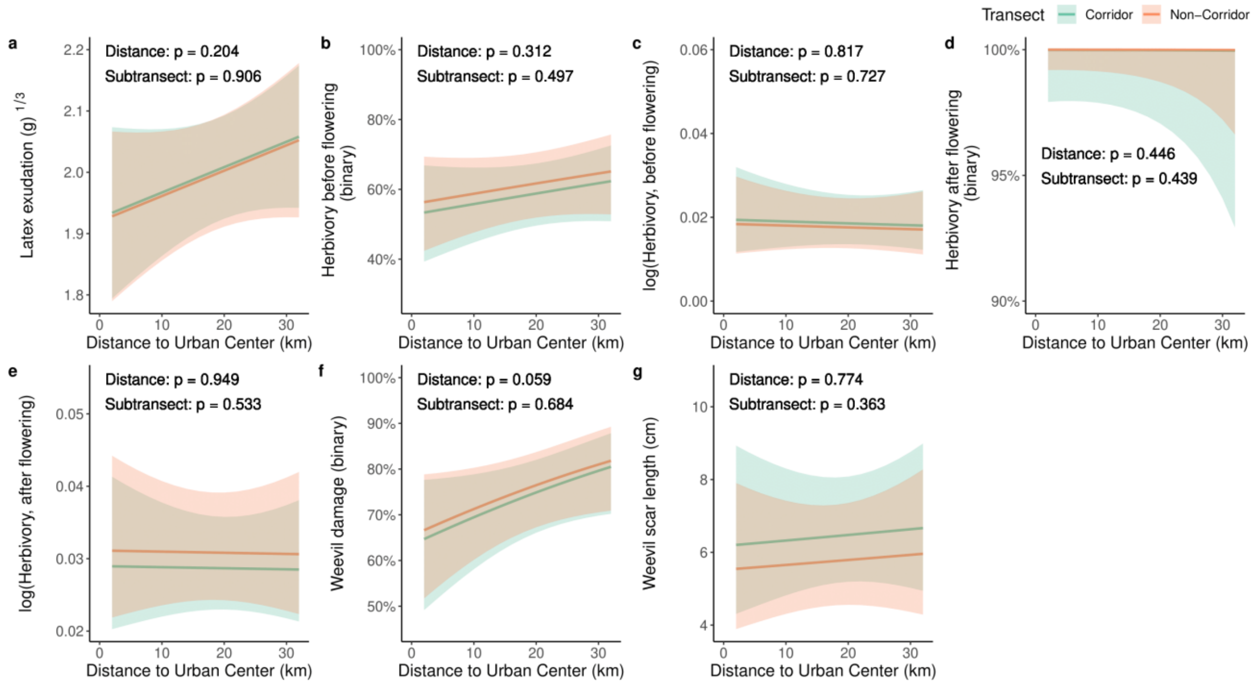


**Supplementary Figure 14.** The effects of urbanization and proximity to a green corridor on plant defense/damage traits when urbanization was quantified by distance from the urban center. Regression lines with a 95% confidence envelope for the mean response, separately for each subtransect, are shown for general and generalized linear mixed effects models.

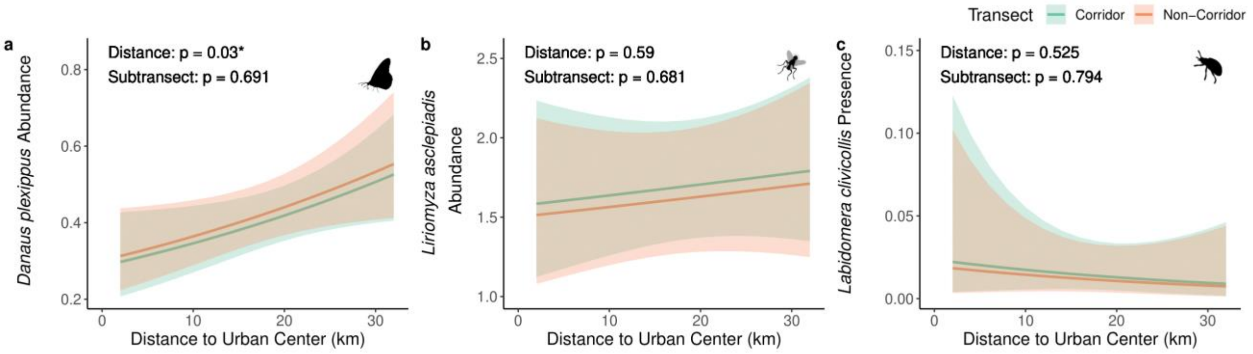


**Supplementary Figure 15.** The effects of urbanization and proximity to a green corridor on herbivore abundance when urbanization was quantified by distance from the urban center. Regression lines with a 95% confidence envelope for the mean response, separately for each subtransect, are shown for generalized linear mixed effects models.

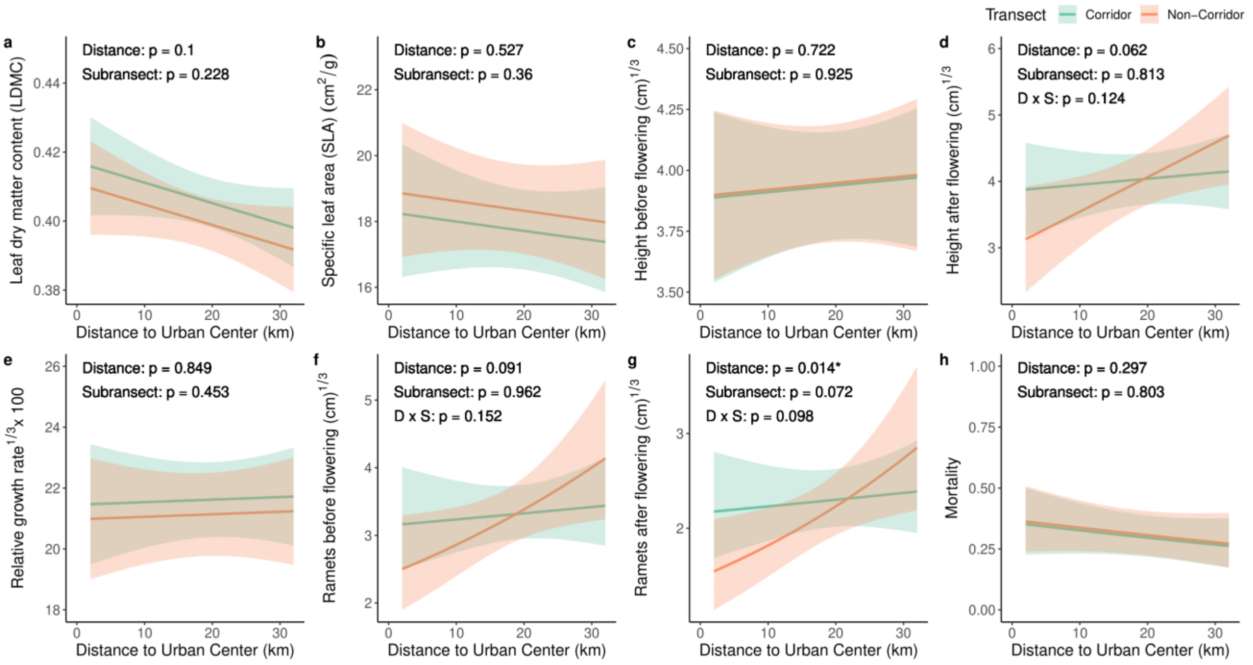


**Supplementary Figure 16.** The effects of urbanization and proximity to a green corridor on plant growth traits when urbanization was quantified by distance from the urban center. Regression lines with a 95% confidence envelope for the mean response, separately for each subtransect, are shown for general and generalized linear mixed effects models.

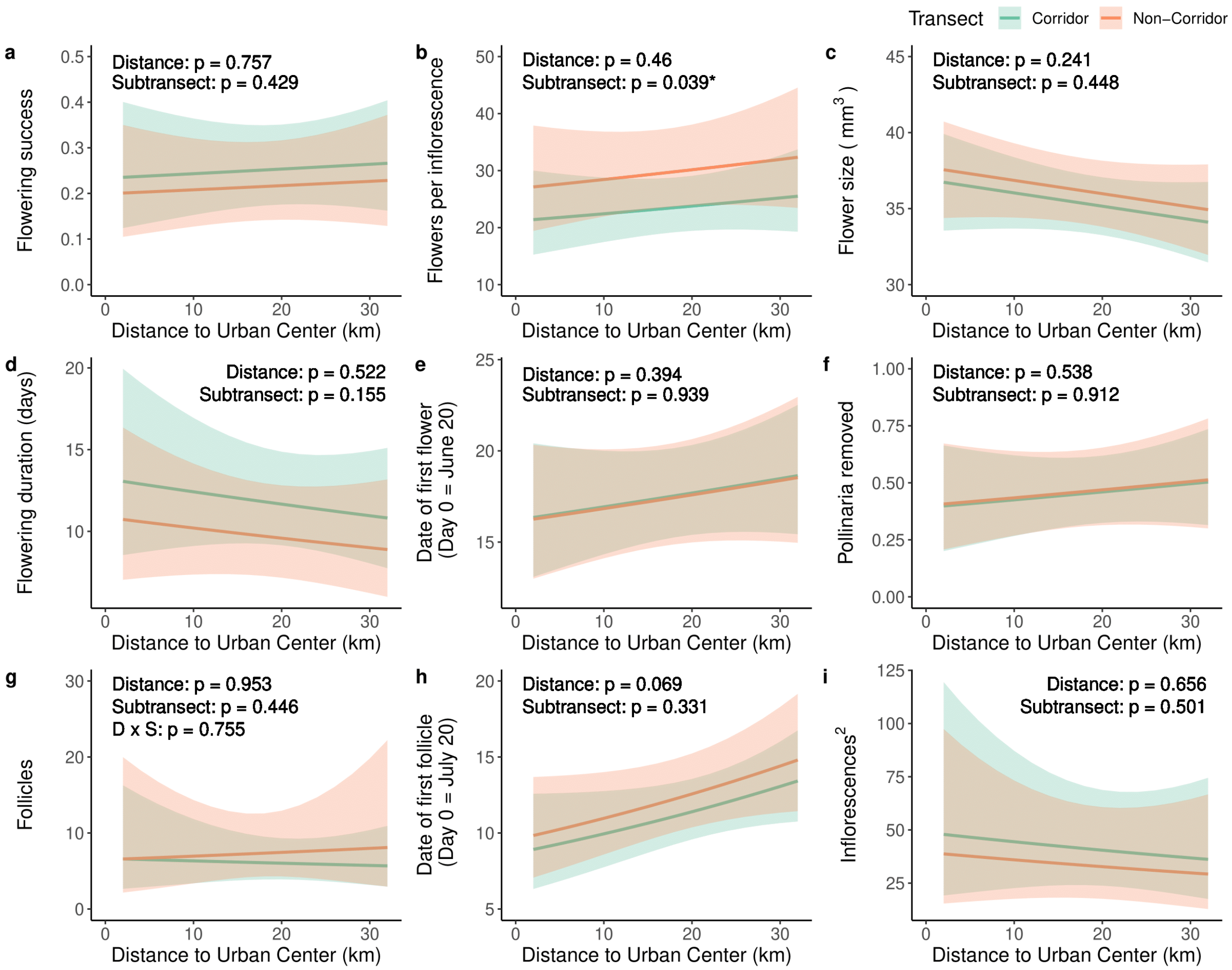


**Supplementary Figure 17.** The effects of urbanization and proximity to a green corridor on plant reproduction traits when urbanization was quantified by distance from the urban center. Regression lines with a 95% confidence envelope for the mean response, separately for each subtransect, are shown for general and generalized linear mixed effects models.

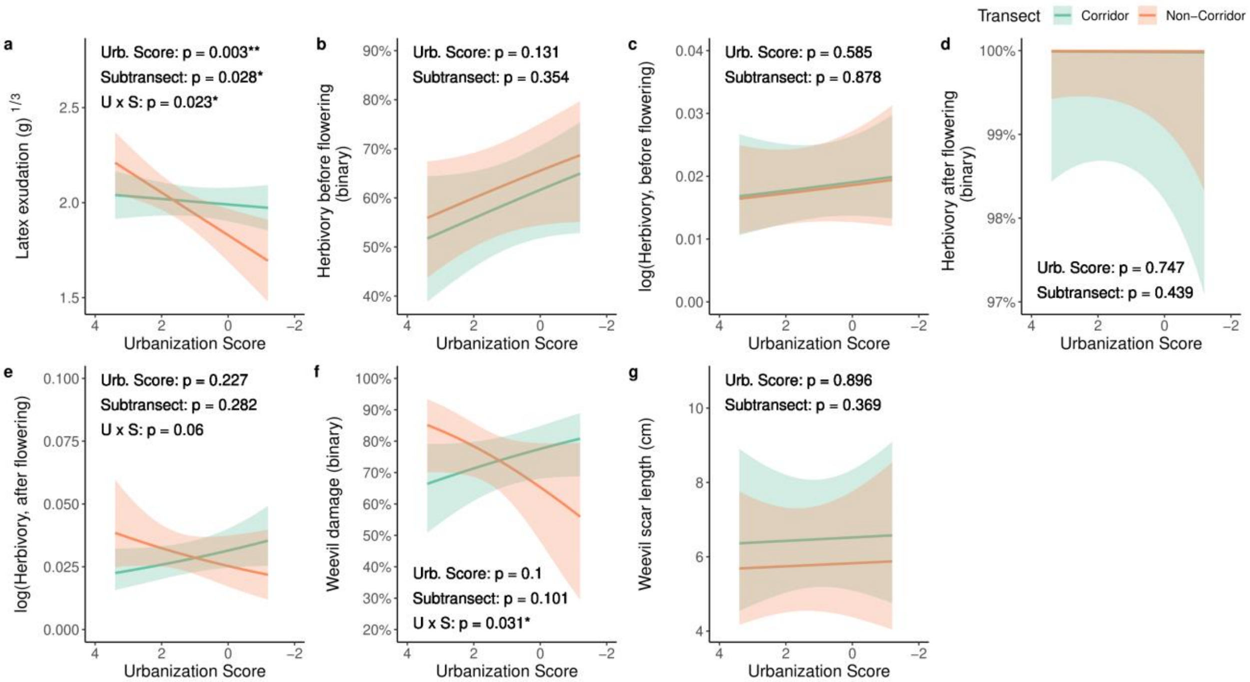


**Supplementary Figure 18.** The effects of urbanization and proximity to a green corridor on plant defense/damage traits when urbanization was quantified by urbanization score. Regression lines with a 95% confidence envelope for the mean response, separately for each subtransect, are shown for general and generalized linear mixed effects models.

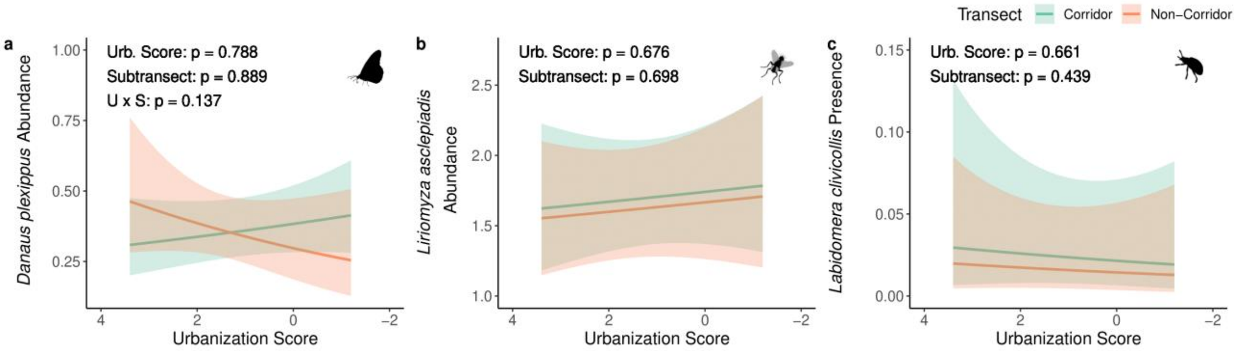


**Supplementary Figure 19.** The effects of urbanization and proximity to a green corridor on herbivore abundance when urbanization was quantified by urbanization score. Regression lines with a 95% confidence envelope for the mean response, separately for each subtransect, are shown for generalized linear mixed effects models.

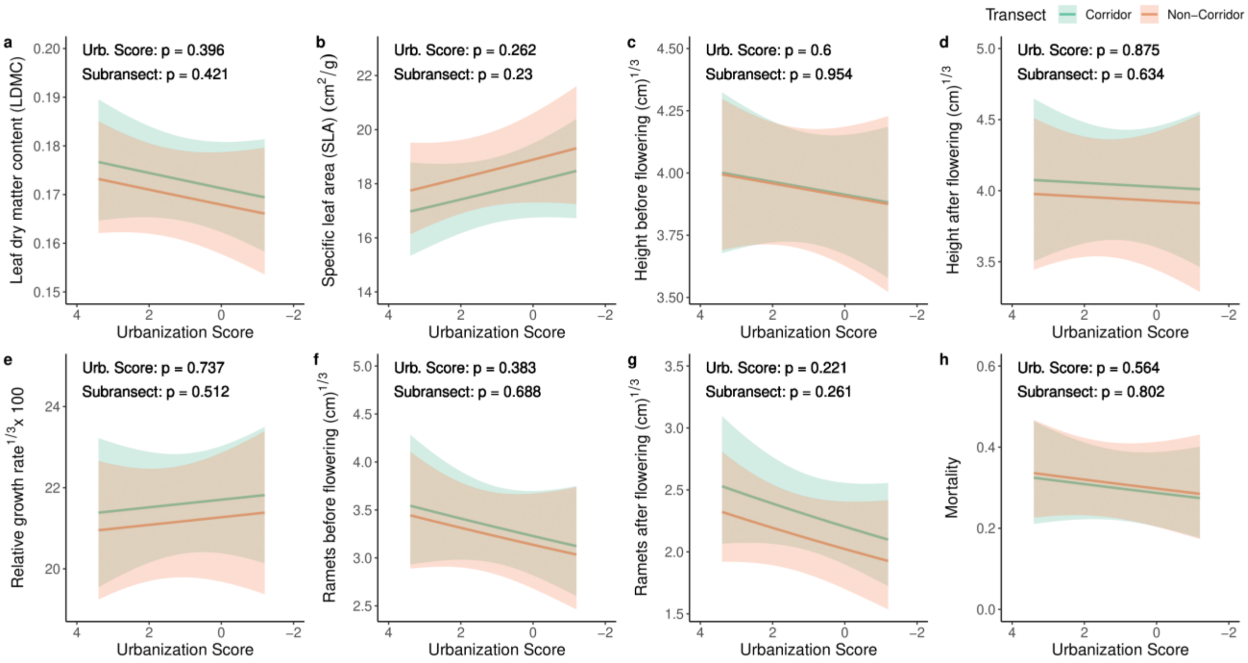


**Supplementary Figure 20.** The effects of urbanization and proximity to a green corridor on plant growth traits when urbanization was quantified by urbanization score. Regression lines with a 95% confidence envelope for the mean response, separately for each subtransect, are shown for general and generalized linear mixed effects models.

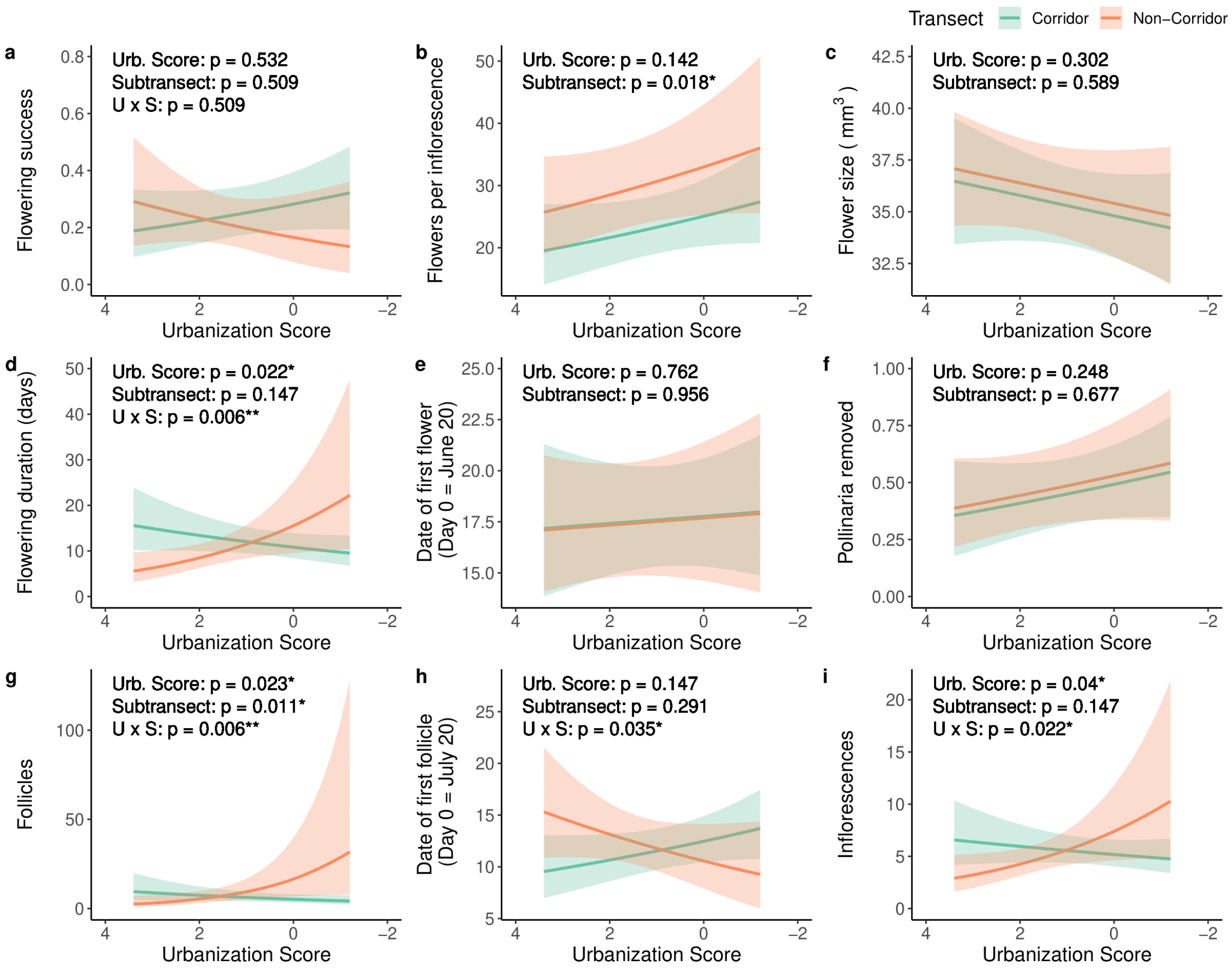


**Supplementary Figure 21.** The effects of urbanization and proximity to a green corridor on plant reproduction traits when urbanization was quantified by urbanization score. Regression lines with a 95% confidence envelope for the mean response, separately for each subtransect, are shown for general and generalized linear mixed effects models.

### Tables

| | Distance | | Urbanization Score | |
|---|---|---|---|---|
| | All Populations | Urban Populations | All Populations | Urban Populations |
| Height before flowering | $x^{1/3}$ | $x^{1/3}$ | $x^{1/3}$ | $x^{1/3}$ |
| Height after flowering | $x^{1/3}$ | $x^{1/3}$ | $x^{1/3}$ | $x^{1/3}$ |
| LDMC | $x^{1/2}$ | $x^{1/2}$ | $x^{1/2}$ | $x^{1/2}$ |
| Mortality | - | - | - | - |
| Ramets before flowering | - | - | - | - |
| Ramets after flowering | - | - | - | - |
| Relative growth rate | $x^{1/3} \times 100$ | $x^{1/3} \times 100$ | $x^{1/3} \times 100$ | $x^{1/3} \times 100$ |
| SLA | log(x + 1) | log(x + 1) | log(x + 1) | log(x + 1) |
| *Danaus plexippus* abundance | - | - | - | - |
| *Labidomera clivicollis* abundance | - | - | - | - |
| *Liriomyza asclepiadis* abundance | - | - | - | - |
| Herbivory before flowering (binary) | - | - | - | - |
| Herbivory before flowering (quantitative) | log(x) | log(x) | log(x) | log(x) |
| Herbivory after flowering (binary) | - | - | - | - |
| Herbivory after flowering (quantitative) | log(x) | log(x) | log(x) | log(x) |
| Latex exudation | $x^{1/3}$ | $x^{1/3}$ | $x^{1/3}$ | $x^{1/3}$ |
| Weevil damage (binary) | - | - | - | - |
| Weevil damage (quantitative) | log(x) | log(x) | log(x) | log(x) |
| Date of first flower | x – 170 | x – 170 | x – 170 | x – 170 |
| Date of first follicle | x – 170 | x – 170 | x – 170 | x – 170 |
| Flower size | - | - | - | - |
| Flowering duration | - | - | - | - |
| Flowering success | - | - | - | - |
| Follicles | - | - | - | - |
| Inflorescences | - | - | $x^2$ | - |
| Flowers per inflorescence | - | - | - | - |
| Pollinaria removed | $x^{1/2}$ | $x^{1/2}$ | $x^{1/2}$ | $x^{1/2}$ |
| Cardenolides | - | - | - | - |

**Supplementary Table 1.** Data transformations used on data to improve normality and homogeneity of variance. “X” represents the transformed response variable. A constant of 170 was subtracted from the dates of first flower and follicle due to model convergence issues.

| | Distance | | Urbanization Score | |
|---|---|---|---|---|
| | All Populations | Urban Populations | All Populations | Urban Populations |
| Height before flowering | - | - | - | - |
| Height after flowering | - | - | - | - |
| LDMC | - | - | - | - |
| Mortality | Binomial | Binomial | Binomial | Binomial |
| Ramets before flowering | Poisson | Poisson | Poisson | Poisson |
| Ramets after flowering | Poisson | Poisson | Poisson | Poisson |
| Relative growth rate | - | - | - | - |
| SLA | - | - | - | - |
| *Danaus plexippus* abundance | Negative binomial | Negative binomial | Negative binomial | Negative binomial |
| *Labidomera clivicollis* abundance | Negative binomial | Negative binomial | Negative binomial | Negative binomial |
| *Liriomyza asclepiadis* abundance | Negative binomial | Negative binomial | Negative binomial | Negative binomial |
| Herbivory before flowering (binary) | Binomial | Binomial | Binomial | Binomial |
| Herbivory before flowering (quantitative) | - | - | - | - |
| Herbivory after flowering (binary) | Binomial | Binomial | Binomial | Binomial |
| Herbivory after flowering (quantitative) | - | - | - | - |
| Latex exudation | - | - | - | - |
| Weevil damage (binary) | Binomial | Binomial | Binomial | Binomial |
| Weevil damage (quantitative) | - | - | - | - |
| Date of first flower | Negative binomial | Negative binomial | Negative binomial | Negative binomial |
| Date of first follicle | Poisson | Poisson | Poisson | Poisson |
| Flower size | - | - | - | - |
| Flowering duration | Negative binomial | Negative binomial | Negative binomial | Negative binomial |
| Flowering success | Binomial | Binomial | Binomial | Binomial |
| Follicles | Negative binomial | Negative binomial | Negative binomial | Negative binomial |
| Inflorescences | Negative binomial | Negative binomial | Negative binomial | Negative binomial |
| Flowers per inflorescence | Negative binomial | Negative binomial | Negative binomial | Negative binomial |
| Pollinaria removed | - | - | - | - |
| Cardenolides | - | - | - | - |

**Supplementary Table 2.** Non-Gaussian probability distributions used on all data.

| Sites | ID | Trait | Urbanization | Predictor | p (Multi-year model) | p (1-year model) |
|---|---|---|---|---|---|---|
| All | 1 | Herbivory before flowering (binary) | Distance | Distance | **0.046** | 0.085 |
| | 2 | Herbivory before flowering (quantitative) | Urb. score | Urb. score | 0.138 | **0.013** |
| | 3 | Flowers per inflorescence | Urb. score | Urb. score | **0.037** | 0.059 |
| Urban | 4 | Herbivory before flowering (quantitative) | Urb. score | Urb. score | **0.041** | 0.585 |
| | 5 | Weevil damage (binary) | Urb. score | U x S | **-** | **0.031** |
| | 6 | Weevil damage (quantitative) | Distance | Subtransect | **0.034** | 0.363 |
| | 7 | Weevil damage (quantitative) | Urb. score | Subtransect | **0.030** | 0.369 |
| | 8 | Date of first follicle | Distance | Distance | **0.011** | 0.069 |
| | 9 | Flowering duration | Distance | Subtransect | **0.013** | 0.155 |
| | 10 | Flowering duration | Urb. Score | Urb. score | 0.396 | **0.022** |
| | 11 | Flowering duration | Urb. Score | U x S | **-** | **0.006** |
| | 12 | Flowers per inflorescence | Distance | Subtransect | 0.071 | **0.039** |
| | 13 | Flowers per inflorescence | Urb. score | Urb. score | **0.032** | 0.142 |
| | 14 | Inflorescences | Urb. score | Subtransect | **0.045** | 0.147 |
| | 15 | Ramets after flowering | Distance | Distance | 0.090 | **0.014** |
| | 16 | Mortality | Distance | Distance | **0.015** | 0.297 |

**Supplementary Table 3.** Discrepancies in type III sums-of-squares ANOVA between models including multiple years of data and the last year of data. Shown are the sites included in the models, model comparison ID, trait, urbanization category, model predictor with a p-value that significantly varied among the multi-year vs. 1-year model, p-value for the multi-year model, and p-value for the 1-year model.

| | Population | | | Family | | |
|---|---|---|---|---|---|---|
| | $\chi^2$ | p | PVE | $\chi^2$ | p | PVE |
| Height before flowering | 0.000 | 0.500 | 0.000 | 1.477 | 0.112 | 3.438 |
| Height after flowering | 0.776 | 0.189 | 1.442 | 5.626 | **0.009** | 6.616 |
| LDMC | 0.000 | 0.380 | 0.407 | 0.000 | 0.500 | 0.000 |
| Mortality | 0.795 | 0.186 | 1.410 | 0.778 | 0.189 | 2.746 |
| Ramets before flowering | 0.032 | 0.429 | 0.000 | 68.887 | **<0.001** | 6.927 |
| Ramets after flowering | 0.121 | 0.364 | 0.324 | 27.650 | **<0.001** | 5.643 |
| Relative growth rate | 0.000 | 0.500 | 0.000 | 0.073 | 0.394 | 1.051 |
| SLA | 0.059 | 0.404 | 0.349 | 0.000 | 0.500 | 0.000 |
| *Danaus plexippus* abundance | 0.070 | 0.396 | 0.354 | 4.088 | **0.021** | 0.000 |
| *Labidomera clivicollis* abundance | 0.000 | 0.500 | 0.675 | 0.000 | 0.500 | 0.000 |
| *Liriomyza asclepiadis* abundance | 1.555 | 0.106 | 2.159 | 1.145 | 0.142 | 2.320 |
| Herbivory before flowering (binary) | 0.000 | 0.500 | 0.000 | 0.001 | 0.490 | 0.317 |
| Herbivory before flowering (quantitative) | 0.000 | 0.500 | 0.000 | 0.000 | 0.500 | 0.000 |
| Herbivory after flowering (binary) | 0.000 | 0.496 | 0.000 | 7.092 | **0.004** | 1.948 |
| Herbivory after flowering (quantitative) | 0.000 | 0.500 | 0.000 | 0.413 | 0.260 | 2.163 |
| Latex exudation | 4.536 | **0.016** | 4.045 | 2.033 | 0.077 | 4.748 |
| Weevil damage (binary) | 0.000 | 0.500 | 0.000 | 3.501 | **0.030** | 5.624 |
| Weevil damage (quantitative) | 1.208 | 0.136 | 2.112 | 1.189 | 0.138 | 4.006 |
| Date of first flower | 0.000 | 0.500 | 0.000 | 0.000 | 0.500 | 9.564 |
| Date of first follicle | 0.000 | 0.500 | 0.000 | 47.484 | **<0.001** | 0.000 |
| Flower size | 1.223 | 0.134 | 7.189 | 0.110 | 0.370 | 4.096 |
| Flowering duration | 0.000 | 0.500 | 0.000 | 0.000 | 0.500 | 0.000 |
| Flowering success | 4.855 | **0.014** | 3.420 | 0.267 | 0.302 | 1.443 |
| Follicles | 0.000 | 0.500 | 0.000 | 0.000 | 0.500 | 0.000 |
| Inflorescences | 3.285 | **0.035** | 7.409 | 0.000 | 0.496 | 0.000 |
| Flowers per inflorescence | 0.833 | 0.180 | 8.369 | 0.000 | 0.500 | 0.000 |
| Pollinaria removed | 0.017 | 0.449 | 1.003 | 3.304 | **0.034** | 27.192 |

**Supplementary Table 4.** Results from general and generalized linear mixed effect models examining the amount of heritable genetic variation within and among populations. All populations were included. Maximum likelihood $\chi^2$ and p-values were obtained from type III sums-of-squares ANOVA performed on random effects.

| | Distance | | | | | | Urbanization Score | | | | | |
|---|---|---|---|---|---|---|---|---|---|---|---|---|
| | Population | | | Family | | | Population | | | Family | | |
| | $\chi^2$ | p | PVE | $\chi^2$ | p | PVE | $\chi^2$ | p | PVE | $\chi^2$ | p | PVE |
| Height before flowering | 0.000 | 0.500 | 0.000 | 1.485 | 0.112 | 3.454 | 0.000 | 0.500 | 0.000 | 1.565 | 0.106 | 3.552 |
| Height after flowering | 0.680 | 0.205 | 1.349 | 5.698 | **0.009** | 6.666 | 0.873 | 0.175 | 1.547 | 5.642 | **0.009** | 6.618 |
| LDMC | 0.000 | 0.424 | 0.254 | 0.000 | 0.500 | 0.000 | 0.157 | 0.346 | 0.537 | 0.000 | 0.500 | 0.000 |
| Mortality | 0.804 | 0.185 | 1.571 | 0.781 | 0.188 | 2.738 | 0.820 | 0.182 | 1.586 | 0.758 | 0.192 | 2.701 |
| Ramets before flowering | 0.001 | 0.485 | 0.000 | 68.905 | **<0.001** | 6.877 | 0.034 | 0.427 | 0.000 | 68.829 | **<0.001** | 7.049 |
| Ramets after flowering | 0.025 | 0.437 | 0.248 | 27.856 | **<0.001** | 5.669 | 0.114 | 0.368 | 0.446 | 27.672 | **<0.001** | 5.650 |
| Relative growth rate | 0.000 | 0.500 | 0.000 | 0.098 | 0.377 | 1.229 | 0.000 | 0.500 | 0.000 | 0.101 | 0.376 | 1.243 |
| SLA | 0.126 | 0.362 | 0.521 | 0.000 | 0.500 | 0.000 | 0.100 | 0.376 | 0.463 | 0.000 | 0.500 | 0.000 |
| *Danaus plexippus* abundance | 0.004 | 0.476 | 0.209 | 4.198 | **0.020** | 0.004 | 0.000 | 0.500 | 0.056 | 4.317 | **0.019** | 0.084 |
| *Labidomera clivicollis* abundance | 0.000 | 0.500 | 0.772 | 0.000 | 0.500 | 0.000 | 0.000 | 0.500 | 0.730 | 0.000 | 0.500 | 0.000 |
| *Liriomyza asclepiadis* abundance | 1.538 | 0.108 | 2.304 | 1.168 | 0.140 | 2.322 | 1.295 | 0.128 | 2.201 | 1.200 | 0.136 | 2.318 |
| Herbivory before flowering (binary) | 0.000 | 0.500 | 0.000 | 0.000 | 0.500 | 0.000 | 0.000 | 0.500 | 0.000 | 0.000 | 0.500 | 0.331 |
| Herbivory before flowering (quantitative) | 0.000 | 0.500 | 0.000 | 0.000 | 0.500 | 0.000 | 0.000 | 0.500 | 0.000 | 0.000 | 0.500 | 0.000 |
| Herbivory after flowering (binary) | 0.000 | 0.494 | 0.000 | 5.815 | **0.008** | 1.937 | 0.001 | 0.486 | 0.000 | 5.699 | **0.009** | 1.894 |
| Herbivory after flowering (quantitative) | 0.000 | 0.500 | 0.000 | 0.495 | 0.241 | 2.370 | 0.000 | 0.500 | 0.000 | 0.482 | 0.244 | 2.348 |
| Latex exudation | 4.628 | **0.016** | 4.141 | 2.052 | 0.076 | 4.770 | 3.010 | **0.042** | 3.217 | 1.907 | 0.084 | 4.631 |
| Weevil damage (binary) | 0.000 | 0.500 | 0.000 | 3.429 | **0.032** | 5.637 | 0.000 | 0.500 | 0.000 | 3.499 | **0.030** | 5.751 |
| Weevil damage (quantitative) | 1.232 | 0.134 | 2.128 | 1.049 | 0.153 | 3.755 | 1.392 | 0.119 | 2.291 | 1.095 | 0.148 | 3.839 |
| Date of first flower | 0.000 | 0.500 | 0.000 | 0.000 | 0.500 | 10.098 | 0.000 | 0.500 | 0.000 | 0.000 | 0.500 | 9.954 |
| Date of first follicle | 0.003 | 0.476 | 0.000 | 45.814 | **<0.001** | 0.000 | 0.000 | 0.500 | 0.000 | 47.014 | **<0.001** | 0.000 |
| Flower size | 1.159 | 0.141 | 7.233 | 0.103 | 0.374 | 3.994 | 0.833 | 0.181 | 6.365 | 0.132 | 0.358 | 4.603 |
| Flowering duration | 0.000 | 0.500 | 0.000 | 0.000 | 0.500 | 0.000 | 0.000 | 0.500 | 0.000 | 0.000 | 0.500 | 0.000 |
| Flowering success | 4.766 | **0.015** | 3.593 | 0.268 | 0.302 | 1.439 | 4.695 | **0.015** | 3.569 | 0.274 | 0.300 | 1.453 |
| Follicles | 0.000 | 0.500 | 0.743 | 0.000 | 0.500 | 0.000 | 0.000 | 0.500 | 0.813 | 0.000 | 0.500 | 0.000 |
| Inflorescences | 3.283 | **0.035** | 8.156 | 0.000 | 0.500 | 0.000 | 3.070 | **0.040** | 7.853 | 0.000 | 0.500 | 0.000 |
| Flowers per inflorescence | 0.203 | 0.326 | 6.489 | 0.000 | 0.500 | 1.507 | 0.060 | 0.403 | 6.237 | 0.000 | 0.500 | 0.838 |
| Pollinaria removed | 0.103 | 0.374 | 2.328 | 2.918 | **0.044** | 25.028 | 0.088 | 0.384 | 2.174 | 2.722 | **0.050** | 24.373 |

**Supplementary Table 5.** Results from general and generalized linear mixed effect models examining the amount of heritable genetic variation associated with urbanization within and among populations. All populations were included. Maximum likelihood $\chi^2$ and p-values were obtained from type III sums-of-squares ANOVA performed on random effects.

| Variable | Predictor | SS | df | F | *p* |
|---|---|---|---|---|---|
| Total Cardenolides | Distance to City Center | 0.049 | 1, 49 | 3.210 | 0.079 |
| | Urbanization Score | 0.002 | 1, 49 | 0.145 | 0.705 |
| Glycosylated Aspecioside | Distance to City Center | 0.037 | 1, 49 | 3.463 | 0.069 |
| | Urbanization Score | 0.002 | 1, 49 | 0.144 | 0.706 |
| Labriformin | Distance to City Center | 0.001 | 1, 49 | 0.482 | 0.491 |
| | Urbanization Score | 0.000 | 1, 49 | 0.385 | 0.538 |
| Cardenolide 17.6 | Distance to City Center | 0.000 | 1, 49 | 0.353 | 0.555 |
| | Urbanization Score | 1.254 | 1, 49 | 3.480 | 0.068 |

**Supplementary Table 6.** Results from general linear models examining the effects of urbanization on cardenolide concentration. All populations were included. Shown are sums of squares (SS), degrees of freedom (df), F statistics, and p-values obtained from type III ANOVA. Cardenolide 17.6 is an unidentified cardenolide with a retention time of 17.6 minutes.

| | | Distance to City Center | | | | Urbanization Score | | | |
|---|---|---|---|---|---|---|---|---|---|
| | | All Populations | | Urban Populations | | All Populations | | Urban Populations | |
| | | Model 1 | | Model 2 | | Model 3 | | Model 4 | |
| **Trait** | **Pseudo-$R^2$ Method** | **$R^2$m** | **$R^2$c** | **$R^2$m** | **$R^2$c** | **$R^2$m** | **$R^2$c** | **$R^2$m** | **$R^2$c** |
| Latex exudation | - | 0.037 | 0.119 | 0.043 | 0.145 | 0.045 | 0.118 | 0.060 | 0.141 |
| Herbivory before flowering (binary) | delta | 0.015 | 0.015 | 0.013 | 0.013 | 0.011 | 0.014 | 0.016 | 0.016 |
| Herbivory before flowering (quantitative) | - | 0.010 | 0.010 | 0.001 | 0.001 | 0.016 | 0.016 | 0.002 | 0.002 |
| Herbivory after flowering (binary) | delta | 0.018 | 0.035 | 0.043 | 0.070 | 0.018 | 0.034 | 0.041 | 0.067 |
| Herbivory after flowering (quantitative) | - | 0.007 | 0.030 | 0.009 | 0.050 | 0.005 | 0.029 | 0.018 | 0.054 |
| Weevil damage (binary) | delta | 0.027 | 0.077 | 0.031 | 0.090 | 0.026 | 0.078 | 0.034 | 0.093 |
| Weevil damage (quantitative) | - | 0.023 | 0.081 | 0.018 | 0.141 | 0.021 | 0.081 | 0.018 | 0.141 |
| Flowering success | delta | 0.038 | 0.088 | 0.037 | 0.109 | 0.038 | 0.088 | 0.042 | 0.114 |
| Flowers per inflorescence | trigamma | 0.000 | 0.001 | 0.000 | 0.001 | 0.001 | 0.001 | 0.000 | 0.001 |
| Flower size | - | 0.019 | 0.129 | 0.034 | 0.197 | 0.021 | 0.128 | 0.030 | 0.208 |
| Flowering duration | trigamma | 0.000 | 0.000 | 0.001 | 0.002 | 0.001 | 0.001 | 0.003 | 0.003 |
| Date of first flower | trigamma | 0.000 | 0.000 | 0.000 | 0.000 | 0.000 | 0.000 | 0.000 | 0.000 |
| Pollinaria removed | - | 0.133 | 0.370 | 0.097 | 0.245 | 0.130 | 0.361 | 0.103 | 0.227 |
| Follicles | trigamma | 0.161 | 0.161 | 0.243 | 0.322 | 0.160 | 0.160 | 0.325 | 0.325 |
| Date of first follicle | trigamma | 0.055 | 0.690 | 0.159 | 0.678 | 0.053 | 0.691 | 0.167 | 0.679 |
| Inflorescences | trigamma | 0.000 | 0.000 | 0.000 | 0.000 | 0.000 | 0.000 | 0.000 | 0.000 |
| *Danaus plexippus* abundance | trigamma | 0.013 | 0.029 | 0.013 | 0.029 | 0.011 | 0.028 | 0.011 | 0.028 |
| *Liriomyza asclepiadis* abundance | trigamma | 0.000 | 0.000 | 0.000 | 0.000 | 0.000 | 0.000 | 0.001 | 0.008 |
| *Labidomera clivicollis* abundance | trigamma | 0.000 | 0.000 | 0.054 | 0.345 | 0.000 | 0.000 | 0.000 | 0.000 |
| LDMC | - | 0.058 | 0.058 | 0.007 | 0.022 | 0.056 | 0.056 | 0.059 | 0.072 |
| SLA | - | 0.065 | 0.071 | 0.068 | 0.108 | 0.065 | 0.071 | 0.070 | 0.106 |
| Height before flowering | - | 0.076 | 0.108 | 0.075 | 0.115 | 0.075 | 0.108 | 0.075 | 0.114 |
| Height after flowering | - | 0.081 | 0.154 | 0.086 | 0.169 | 0.079 | 0.154 | 0.074 | 0.164 |
| Relative growth rate | - | 0.014 | 0.026 | 0.021 | 0.031 | 0.014 | 0.027 | 0.021 | 0.030 |
| Ramets before flowering | trigamma | 0.080 | 0.269 | 0.094 | 0.297 | 0.078 | 0.270 | 0.081 | 0.290 |
| Ramets after flowering | trigamma | 0.102 | 0.204 | 0.121 | 0.224 | 0.100 | 0.204 | 0.111 | 0.217 |
| Mortality | delta | 0.035 | 0.075 | 0.040 | 0.123 | 0.035 | 0.075 | 0.039 | 0.122 |

**Supplementary Table 7.** Marginal and conditional $R^2$ values from general and generalized linear mixed effect models examining the effects of urbanization and a green corridor on all phenotypic traits. Shown are traits, method for deriving the observation-level variance for generalized linear mixed effect models when calculated as pseudo-$R^2$, and marginal and conditional $R^2$ values when urbanization was quantified by distance from the urban center (Models 1-2) and urbanization score (Models 3-4), and when all populations were included (Models 1 & 3) and only urban sites were included (Models 2 & 4).

| **Variable** | **Urbanization** | **$R^2$** | **$R^2_{adj}$** |
|---|---|---|---|
| Total Cardenolides | Distance to City Center | 0.061 | 0.042 |
| | Urbanization Score | 0.003 | -0.017 |
| Glycosylated Aspecioside | Distance to City Center | 0.066 | 0.047 |
| | Urbanization Score | 0.003 | -0.017 |
| Labriformin | Distance to City Center | 0.010 | -0.010 |
| | Urbanization Score | 0.008 | -0.012 |
| Cardenolide 17.6 | Distance to City Center | 0.007 | -0.013 |
| | Urbanization Score | 0.066 | 0.047 |

**Supplementary Table 8.** Standard and adjusted $R^2$ values from general linear models examining the effects of urbanization on cardenolide concentrations. All populations were included. Cardenolide 17.6 is an unidentified cardenolide with a retention time of 17.6 minutes.

| | Distance | | | | | | Urbanization Score | | | | | |
|---|---|---|---|---|---|---|---|---|---|---|---|---|
| | Population | | | Family | | | Population | | | Family | | |
| | $\chi^2$ | p | PVE | $\chi^2$ | p | PVE | $\chi^2$ | p | PVE | $\chi^2$ | p | PVE |
| Height before flowering | 0.000 | 0.500 | 0.000 | 1.681 | 0.098 | 4.617 | 0.000 | 0.500 | 0.000 | 1.575 | 0.104 | 4.475 |
| Height after flowering | 0.000 | 0.500 | 0.000 | 7.154 | **0.004** | 9.027 | 0.050 | 0.411 | 0.485 | 7.029 | **0.004** | 9.265 |
| LDMC | 0.050 | 0.412 | 0.413 | 0.011 | 0.459 | 0.439 | 0.211 | 0.323 | 0.888 | 0.013 | 0.454 | 0.492 |
| Mortality | 0.000 | 0.500 | 1.067 | 3.952 | **0.024** | 7.163 | 0.000 | 0.500 | 0.923 | 3.965 | **0.023** | 7.079 |
| Ramets before flowering | 0.000 | 0.500 | 0.165 | 52.455 | **<0.001** | 7.884 | 0.000 | 0.500 | 0.473 | 52.334 | **<0.001** | 7.827 |
| Ramets after flowering | 0.000 | 0.500 | 0.000 | 22.590 | **<0.001** | 5.826 | 0.000 | 0.500 | 0.000 | 22.903 | **<0.001** | 5.999 |
| Relative growth rate | 0.483 | 0.244 | 1.389 | 0.000 | 0.500 | 0.000 | 0.237 | 0.314 | 0.973 | 0.000 | 0.500 | 0.000 |
| SLA | 2.440 | 0.059 | 3.728 | 0.037 | 0.424 | 0.785 | 2.258 | 0.066 | 3.486 | 0.006 | 0.469 | 0.319 |
| *Danaus plexippus* abundance | 0.000 | 0.500 | 0.000 | 0.000 | 0.500 | 0.000 | 0.017 | 0.448 | 0.437 | 0.044 | 0.418 | 0.000 |
| *Labidomera clivicollis* abundance | 0.000 | 0.500 | 0.800 | 0.000 | 0.500 | 0.000 | 0.000 | 0.500 | 0.000 | 0.000 | 0.500 | 0.000 |
| *Liriomyza asclepiadis* abundance | 0.034 | 0.428 | 1.673 | 3.365 | **0.034** | 4.881 | 0.055 | 0.407 | 1.583 | 3.181 | **0.038** | 4.849 |
| Herbivory before flowering (binary) | 0.000 | 0.500 | 0.000 | 0.000 | 0.500 | 0.000 | 0.000 | 0.500 | 0.000 | 0.000 | 0.500 | 0.000 |
| Herbivory before flowering (quantitative) | 0.000 | 0.500 | 0.000 | 0.000 | 0.500 | 0.000 | 0.000 | 0.500 | 0.000 | 0.000 | 0.500 | 0.000 |
| Herbivory after flowering (binary) | 0.001 | 0.490 | 0.000 | 0.315 | 0.288 | 3.729 | 0.001 | 0.488 | 0.000 | 0.280 | 0.298 | 3.809 |
| Herbivory after flowering (quantitative) | 0.000 | 0.500 | 0.000 | 1.076 | 0.150 | 4.410 | 0.000 | 0.500 | 0.000 | 0.743 | 0.194 | 3.628 |
| Latex exudation | 2.000 | 0.055 | 3.946 | 3.074 | **0.040** | 7.114 | 0.504 | 0.239 | 1.722 | 3.228 | **0.036** | 7.465 |
| Weevil damage (binary) | 0.000 | 0.500 | 0.000 | 2.741 | **0.049** | 6.715 | 0.000 | 0.500 | 0.000 | 2.614 | 0.053 | 6.614 |
| Weevil damage (quantitative) | 2.144 | 0.072 | 4.356 | 3.595 | **0.029** | 8.479 | 1.603 | 0.102 | 3.912 | 3.715 | **0.027** | 8.672 |
| Date of first flower | 0.000 | 0.500 | 0.000 | 0.000 | 0.500 | 12.682 | 0.000 | 0.500 | 0.000 | 0.000 | 0.500 | 10.791 |
| Date of first follicle | 0.000 | 0.500 | 0.000 | 29.669 | **<0.001** | 0.000 | 0.000 | 0.500 | 0.000 | 30.383 | **<0.001** | 0.000 |
| Flower size | 1.331 | 0.124 | 10.569 | 0.319 | 0.286 | 8.428 | 0.845 | 0.179 | 9.030 | 0.414 | 0.260 | 9.935 |
| Flowering duration | 0.000 | 0.500 | 0.769 | 0.000 | 0.500 | 0.000 | 0.000 | 0.500 | 0.000 | 0.000 | 0.500 | 0.000 |
| Flowering success | 0.115 | 0.368 | 2.045 | 2.696 | 0.051 | 5.187 | 0.025 | 0.437 | 1.459 | 2.814 | **0.046** | 5.308 |
| Follicles | 0.000 | 0.500 | 14.430 | 0.000 | 0.500 | 11.954 | 0.000 | 0.500 | 5.764 | 0.000 | 0.500 | 11.434 |
| Inflorescences | 1.325 | 0.125 | 12.878 | 9.892 | **0.001** | 0.000 | 0.930 | 0.168 | 7.011 | 9.238 | **0.001** | 0.000 |
| Flowers per inflorescence | 0.359 | 0.274 | 11.915 | 0.000 | 0.500 | 1.850 | 0.000 | 0.500 | 9.792 | 0.000 | 0.500 | 1.128 |
| Pollinaria removed | 0.145 | 0.352 | 3.139 | 0.628 | 0.214 | 14.544 | 0.035 | 0.426 | 1.477 | 0.345 | 0.278 | 11.331 |

**Supplementary Table 9.** Results from general and generalized linear mixed effect models examining the amount of heritable genetic variation associated with urbanization and proximity to a green corridor within and among populations. Only urban populations were included. Maximum likelihood $\chi^2$ and p-values were obtained from type III sums-of-squares ANOVA performed on random effects.

| | Urbanization Score | | Subtransect | | U x S | |
|---|---|---|---|---|---|---|
| | $\chi^2$ | p | $\chi^2$ | p | $\chi^2$ | p |
| Latex exudation | 9.011 | **0.003**** | 4.840 | **0.028*** | 5.164 | **0.023*** |
| Herbivory before flowering (binary) | 2.286 | 0.131 | 0.860 | 0.354 | | |
| Herbivory before flowering (quantitative) | 0.298 | 0.585 | 0.023 | 0.878 | | |
| Herbivory after flowering (binary) | 0.104 | 0.747 | 0.600 | 0.439 | | |
| Herbivory after flowering (quantitative) | 1.457 | 0.227 | 1.155 | 0.282 | 3.528 | 0.060 |
| Weevil damage (binary) | 2.701 | 0.100 | 2.684 | 0.101 | 4.667 | **0.031*** |
| Weevil damage (quantitative) | 0.017 | 0.896 | 0.806 | 0.369 | | |
| Flowering success | 0.391 | 0.532 | 0.435 | 0.509 | 2.088 | 0.148 |
| Flowers per inflorescence | 2.158 | 0.142 | 5.572 | **0.018*** | | |
| Flower size | 1.066 | 0.302 | 0.292 | 0.589 | | |
| Flowering duration | 5.274 | **0.022*** | 2.108 | 0.147 | 7.614 | **0.006**** |
| Date of first flower | 0.092 | 0.762 | 0.003 | 0.956 | | |
| Follicles | 5.185 | **0.023*** | 6.494 | **0.011*** | 7.454 | **0.006**** |
| Date of first follicle | 2.108 | 0.147 | 1.113 | 0.291 | 4.467 | **0.035*** |
| Inflorescences | 4.233 | **0.040*** | 2.101 | 0.147 | 5.252 | **0.022*** |
| Pollinaria removed | 1.336 | 0.248 | 0.173 | 0.677 | | |
| *Danaus plexippus* abundance | 0.072 | 0.788 | 0.019 | 0.889 | 2.213 | 0.137 |
| *Liriomyza asclepiadis* abundance | 0.175 | 0.676 | 0.150 | 0.698 | | |
| *Labidomera clivicollis* abundance | 0.193 | 0.661 | 0.600 | 0.439 | | |
| LDMC | 0.720 | 0.396 | 0.646 | 0.421 | | |
| SLA | 1.260 | 0.262 | 1.438 | 0.230 | | |
| Height before flowering | 0.275 | 0.600 | 0.003 | 0.954 | | |
| Height after flowering | 0.025 | 0.875 | 0.227 | 0.634 | | |
| Relative growth rate | 0.113 | 0.737 | 0.431 | 0.512 | | |
| Ramets before flowering | 0.761 | 0.383 | 0.161 | 0.688 | | |
| Ramets after flowering | 1.499 | 0.221 | 1.263 | 0.261 | | |
| Mortality | 0.332 | 0.564 | 0.063 | 0.802 | | |

**Supplementary Table 10.** Results from general and generalized linear mixed effect models examining the effects of urbanization and proximity to a green corridor on all phenotypic traits. Urbanization was quantified via urbanization score and only urban populations were included. Shown are maximum likelihood $\chi^2$ and p-values obtained from type III sums-of-squares ANOVA. Though not shown, block was included as a fixed effect and often explained significant variation in the common garden.
